# A conserved odorant receptor underpins borneol-mediated repellency in culicine mosquitoes

**DOI:** 10.1101/2023.08.01.548337

**Authors:** Yuri Vainer, Yinliang Wang, Robert M. Huff, Dor Perets, Evyatar Sar-Shalom, Esther Yakir, Majid Ghaninia, Iliano V. Coutinho-Abreu Gomes, Carlos Ruiz, Dhivya Rajamanickam, A. Warburg, Omar S. Akbari, Philippos A. Papathanos, R. Ignell, Jeffrey A. Riffell, R. Jason Pitts, Jonathan D. Bohbot

**Affiliations:** Department of Biology, Baylor University, USA; Department of Entomology, The Hebrew University of Jerusalem, Israel; Division of Biological Sciences, University of California, USA; Department of Microbiology and Molecular Genetics, The Hebrew University of Jerusalem, Israel; Department of Plant Protection Biology, Swedish University of Agricultural Sciences, Sweden; Department of Biology, University of Washington, USA; Northeast Normal University, China

**Keywords:** mosquitoes, monoterpenoids, borneol, camphor, repellent, odorant receptor

## Abstract

The use of essential oils derived from the camphor tree to repel mosquitoes is an ancient practice that originated in Southeast Asia and gradually spread to China and across Europe via the Maritime Silk Road. The olfactory mechanisms by which these oils elicit avoidance behavior are unclear. Here we show that plant bicyclic monoterpenoids and borneol specifically activate a neural pathway that originates in the orphan olfactory receptor neuron of the capitate peg sensillum in the maxillary palp, and projects to the mediodorsal glomerulus 3 in the antennal lobe. This neuron co-locates with two olfactory receptor neurons tuned to carbon dioxide and octenol that mediate human-host detection. We also confirm that borneol elicits repellency against human-seeking female mosquitoes. Understanding the functional role of the mosquito maxillary palp is essential to investigating olfactory signal integration and host-selection behavior.

## Main

The use of plants to ward off insects has been a human practice since prehistorical times ^1^ and is still used in many parts of the world ^2,3^. Plant-based essential oils (EOs), including lemon eucalyptus leaf oil ^4^, citronella oil ^5^, and coconut oil ^6^ exhibit different degrees of mosquito repellency due to the presence of pyrethrins ^7^, phenol derivatives, and terpenoids ^8^. The latter includes oxygen-containing compounds with open-chain hydrocarbons, such as linalool, citronellol, geraniol, and bicyclic derivatives such as cineole, fenchone, camphor and borneol (Fig. 1a). The EO of the camphor tree *Cinnamomum camphora* is composed of over 70 volatile organic compounds (VOCs), most of which are oxygenated monoterpenes dominated by camphor and other related compounds ^9^. Camphor and borneol extracts are believed to have originated from the camphor tree in the island of Borneo (Fig. 1a), where they were initially traded with China and then introduced to the West due to their therapeutic, refreshing, and repellent effects against mosquitoes ^10^. How these terpenoids molecules exert repelling effects against mosquitoes is not well-understood but is likely mediated by their olfactory system.

**Figure 1.**
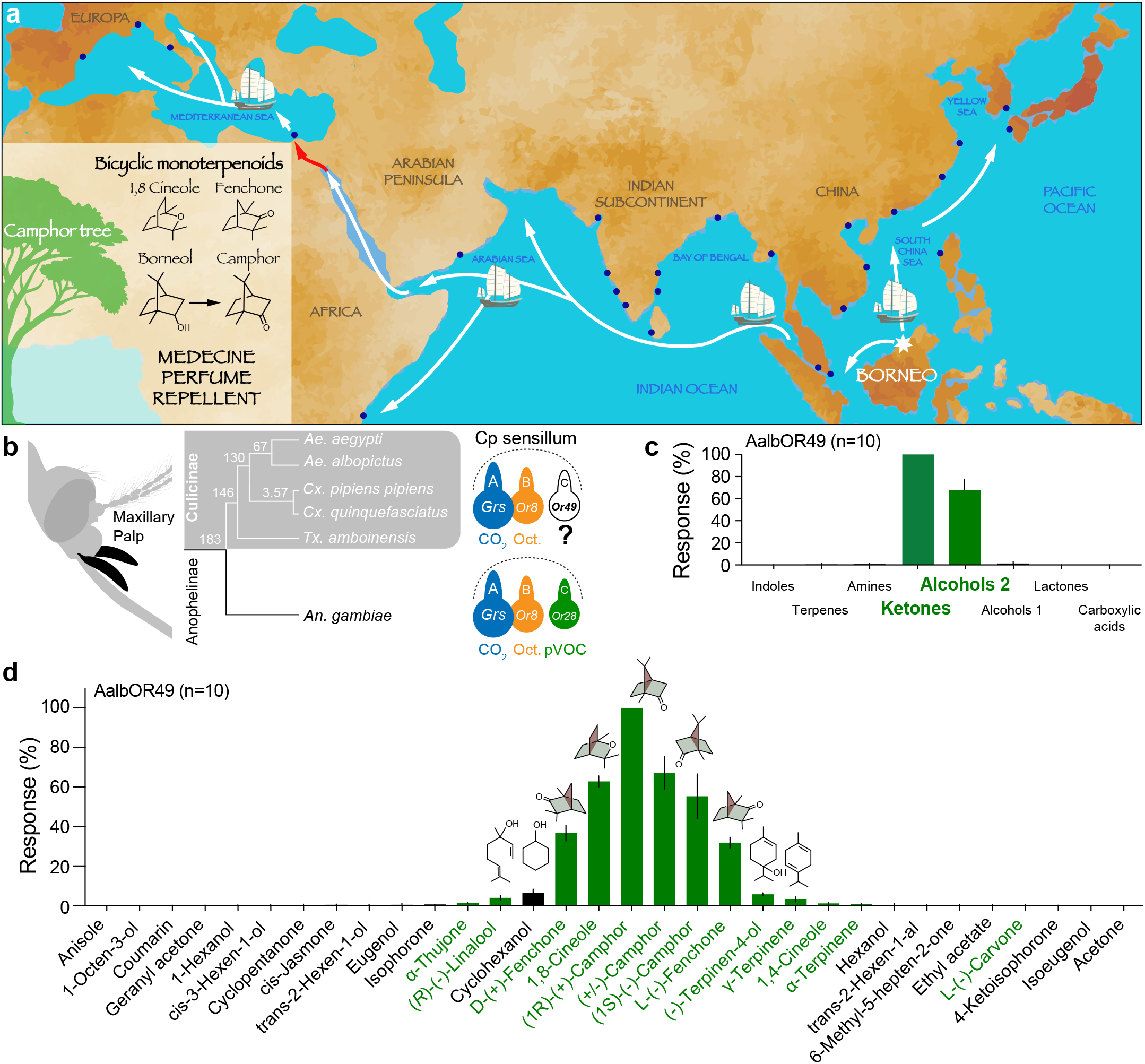
Odorant receptor 49 is activated by bicyclic monoterpenoids. **a**, Borneol and camphor oils from the camphor tree *Cinnamomum camphora* originated from the island of Borneo and were traded with China and the rest of the Western world through the maritime silk road during the classic age due to their medicinal and repellent properties. The world map is a modified illustration Vecteezy.com (free license) where blue dots indicate major trading ports. **b**, The capitate peg sensillum (cp) on the 4th segment of the mosquito maxillary palps houses three neurons, including the CO_2_ capitate-peg neuron (cpA) expressing three gustatory receptors, the 1-octen-3-ol-sensitive neuron (cpB), and the orphan neuron (cpC). **c**, The response profile of *Aedes albopictus* OR49 (AalbOR49) to odorant mixtures (100 μM) belonging to a variety of chemical classes highlights the activity of ketones. **d**, Bicyclic monoterpenoids, including camphor and fenchone are the most efficacious activators of AalbOR49 (labeled in green). Statistical differences were evaluated by ANOVA followed by a Kruskal-Wallis multiple comparisons test. ns, non-significant, *p < 0.0332, **p < 0.0021, ***p < 0.0002 and ****p < 0.0001. Data indicate the means ± SEM.

The capitate-peg (cp) sensillum located on the mosquito maxillary palp comprises three olfactory neurons, each distinguishable by size, olfactory receptor gene expression profile, and odor response characteristics (Fig. 1b). In both culicine and anopheline mosquitoes, the largest olfactory sensory neuron (cpA) expresses three gustatory receptors (*Grs*) that specifically detect CO_2_ ^11,12^. The medium-sized cpB neuron of *Anopheles gambiae* and *Aedes aegypti* expresses the 1-octen-3-ol odorant receptor *Or8* and its co-receptor Orco ^13–15^. Both CO_2_ and 1-octen-3-ol elicit attraction and signal the presence of animal hosts in anopheline ^16^ and culicine ^17^ mosquitoes. This cellular and functional organization have remained remarkably conserved over 180 million years of mosquito evolution (Fig. 1b).

In anophelines, the small cpC neuron expresses the *Or28* gene ^13^, which responds to plant volatile organic compounds (pVOC) ^18^. In culicines, the cpC neuron expresses the *Or49* gene ^19^, which is unrelated to *AgamOr28* (Fig. 1b). Using a pharmacological approach, we expressed *Or49* in a heterologous expression system and exposed it to odorant mixtures, EOs and individual volatile odorant compounds. Our findings provide strong evidence that the OR49 receptor and the cpC neuron respond to plant-derived bicyclic monoterpenoids with a marked selectivity towards borneol, traditionally used as a mosquito repellent. We observed that a loss-of-function mutation in the *Or49* gene leads to a lack of electrophysiological responses when stimulated by borneol. Furthermore, we show that borneol elicits selective activation of the MD3 glomerulus in the antennal lobe of *Ae. aegypti*, indicating that all 3 cp neurons project to the same brain area. Confirming our pharmacological and electrophysiological results, our behavioral study reveals that borneol induces repellency in human-seeking female mosquitoes. our findings lay the groundwork to understand the detailed molecular and neural basis shaping olfactory integration and governing host selection in mosquitoes.

### Odorant receptor 49 is an evolutionary conserved borneol receptor

The odorant receptor 49 gene^18,19^ is conserved in *Aedes albopictus, Culex quinquefasciatus, Toxorhynchites amboinensis* and *Anopheles gambiae* (Supplementary Fig. 1). Phylogenetic analyses support previous studies ^19,20^ describing the OR49 family as a Culicinae-specific group (Supplementary Fig. 1c–d) that includes the more distantly related anopheline OR48/49 group. The *Or49* genes are located on chromosome 2 in both *Aedes* and *Anopheles* mosquitoes (Supplementary Fig. 1e) and exhibit conserved syntenic relationships with neighbouring genes (Supplementary Fig. 1f).

As a continuation of our work on the functional identity of the cp sensillum in culicine mosquitoes ^19,21^, we investigated the receptive field of *Ae. albopictus* OR49 (AalbOR49) using two-electrode voltage clamp recordings (Fig. 1c–d) and a panel of 81 odorants representing a variety of chemical classes (Supplementary Table 1). Oocytes expressing functional receptor complexes were strongly activated by blends containing ketonic compounds (Fig. 1c). Individual testing of the single compounds from the ketone blend revealed the efficacy of bicyclic monoterpenoids, with camphor emerging as the most efficacious member followed by cineole (also called eucalyptol) and fenchones (Fig. 1d). Subsequently, we screened OR49 from the nectar-feeding *Tx. amboinensis* (*TambOR49*) with 8 *Cannabis* EOs (Fig. 2a, Supplementary Table 1) containing 42 plant VOCs dominated by terpenes (Supplementary Fig. 2). The three most active mixtures were Pineapple Haze, OG Kush, Adom #9 followed by Master Kush with a much lower efficacy (Fig. 2a), all evoking consistent current responses (Supplementary Fig. 3). Successive fractionations of these EOs down to 6 sub-mixtures and 4 single compounds were then administered of which the racemic borneol showed the highest potency (Fig. 2b), consistent with the borneol content in the original *Cannabis* mixtures (Fig. 2a, Supplementary Fig. 4a) and sub-mixtures (Supplementary Fig. 4b).

**Figure 2.**
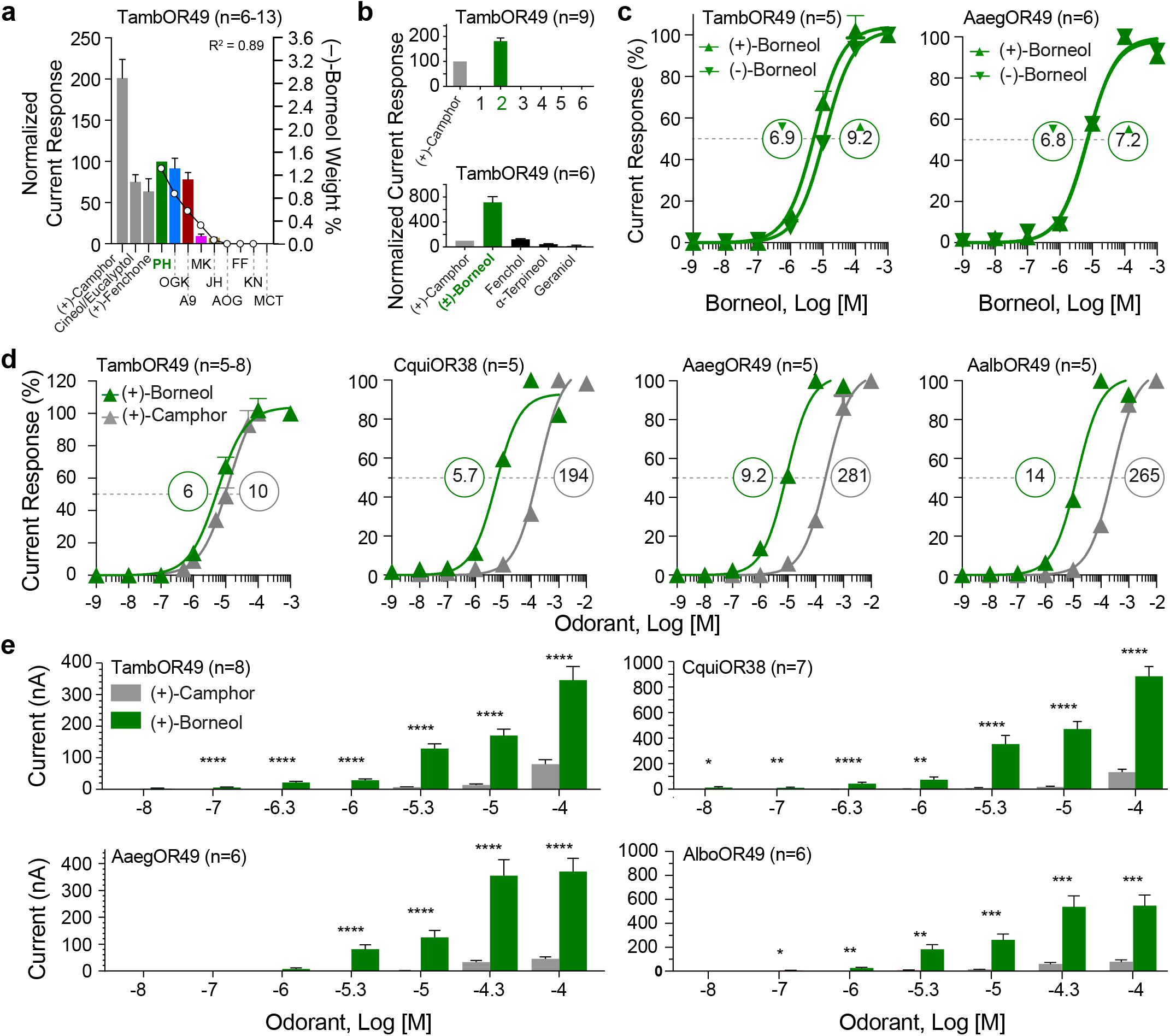
Odorant receptor 49 is a borneol receptor. **a**, Pineapple Haze (PH), OG Kush (OGK), Adom9 (A9), and Master Kush (MK) elicited the highest currents from *Toxorhynchites amboinensis* OR49 (TambOR49). Borneol content is indicated on the right Y-axis. A correlation between the OR49 response and the borneol content in each Cannabis EO was observed (R^2^ = 0.89). **b**, Response of TambOR49 to 6 *Cannabis* sub-mixtures and the constituents from sub-mixture 2. **c**, Concentration-response relationships of TambOR49 and AaegOR49 to the two borneol enantiomers. **d**, Concentration-response relationships of TambOR49, *Culex quinquefasciatus* OR38 (CquiOR38), *Ae. aegypti* OR49 (AaegOR49) and AalbOR49 in response to (+)-borneol and (+)-camphor. Effective concentrations at 50% of the maximal response (EC_50_) are circled. **e**) Pairwise comparisons of the current responses of all four mosquito species OR49s elicited by increasing concentrations of (+)-borneol and (+)-camphor. Statistical differences were evaluated by multiple t-tests. *p < 0.0332, **p < 0.0021, ***p < 0.0002 and ****p < 0.0001. Data indicate the means ± SEM. Representative current traces can be found in Supplementary Fig. 3.

In *An. gambiae*, the cpC neuron expresses OR28 (Figure 1a), which is activated by plant VOCs, with acetophenone and 2,4,5-trimethylthiazole being the most effective ligands in the *Xenopus* oocyte expression system ^13,18^. The EC_50_ values of acetophenone, α-pinene, and α-terpineol were in the low millimolar range indicating that more potent ligands remain to be identified (Supplementary Fig. 5). Borneol did not elicit any response at the tested concentrations suggesting that AgamOR28 is tuned to VOCs belonging to a different chemical class.

TambOR49 and AaegOR49 did not exhibit enantioselectivity towards the (+) and (–)-borneol (Figure 2c) as indicated by their EC_50_ values in the one-digit micromolar range. We established concentration-response relationships between TambOR49, *Cx. quinquefasciatus* OR38 (CquiOR49), *Ae. aegypti* OR49 (AaegOR49), and *Ae. allbopictus* OR49 (AalbOR49), for the two most potent ligands, (+)-borneol and (+)-camphor (Fig. 2d). (+)-Borneol was 19-32 times more potent than (+)-camphor in all cases except for TambOR49 in which both compounds were equally active in the low micromolar range. In all four examined culicine species, including the nectar-feeding *Tx. amboinensis*, (+)-borneol elicited significantly higher responses than (+)-camphor across all concentrations (Fig. 2e, Supplementary Fig. 3). This data indicates that OR49 is a selective borneol receptor.

### The *Or49* gene confers sensitivity to borneol

To investigate the olfactory effect of camphor and borneol *in vivo*, we conducted electropalpogram (EPG) recordings on three culicine (*Ae. albopictus, Ae. aegypti, Cx. pipiens*) and one anopheline (*An. gambiae*) species. 1-Octen-3-ol, which served as a positive control, induced significant responses in all mosquitoes tested. Camphor and borneol elicited consistent palp responses in Culicinae mosquitoes, including *Ae. albopictus, Ae. aegypti*, and *Cx. pipiens* but not in *An. gambiae* (Fig. 3a). Camphor and borneol evoked lower responses than 1-octen-3-ol in the culicine palps, which is consistent with the smaller size of the cpC neuron in comparison to the cpB neuron. In *Ae. albopictus*, (–)-borneol was more active than (+)-camphor, while in *Ae. aegypti* both borneol enantiomers elicited significantly greater EPG responses than camphor enantiomers. In *Cx. pipiens*, borneol and camphor elicited comparable EPG responses. We established dose-response relationships with (+)-camphor and (+)-borneol in *Ae. albopictus, Ae. aegypti*, and *Cx. pipiens* (Fig. 3b). Among the three culicine species, *Cx. pipiens* exhibited the largest responses in response to (+)-camphor. However, we did not find any significant statistical differences in activity between camphor and borneol in the tested species.

**Figure 3.**
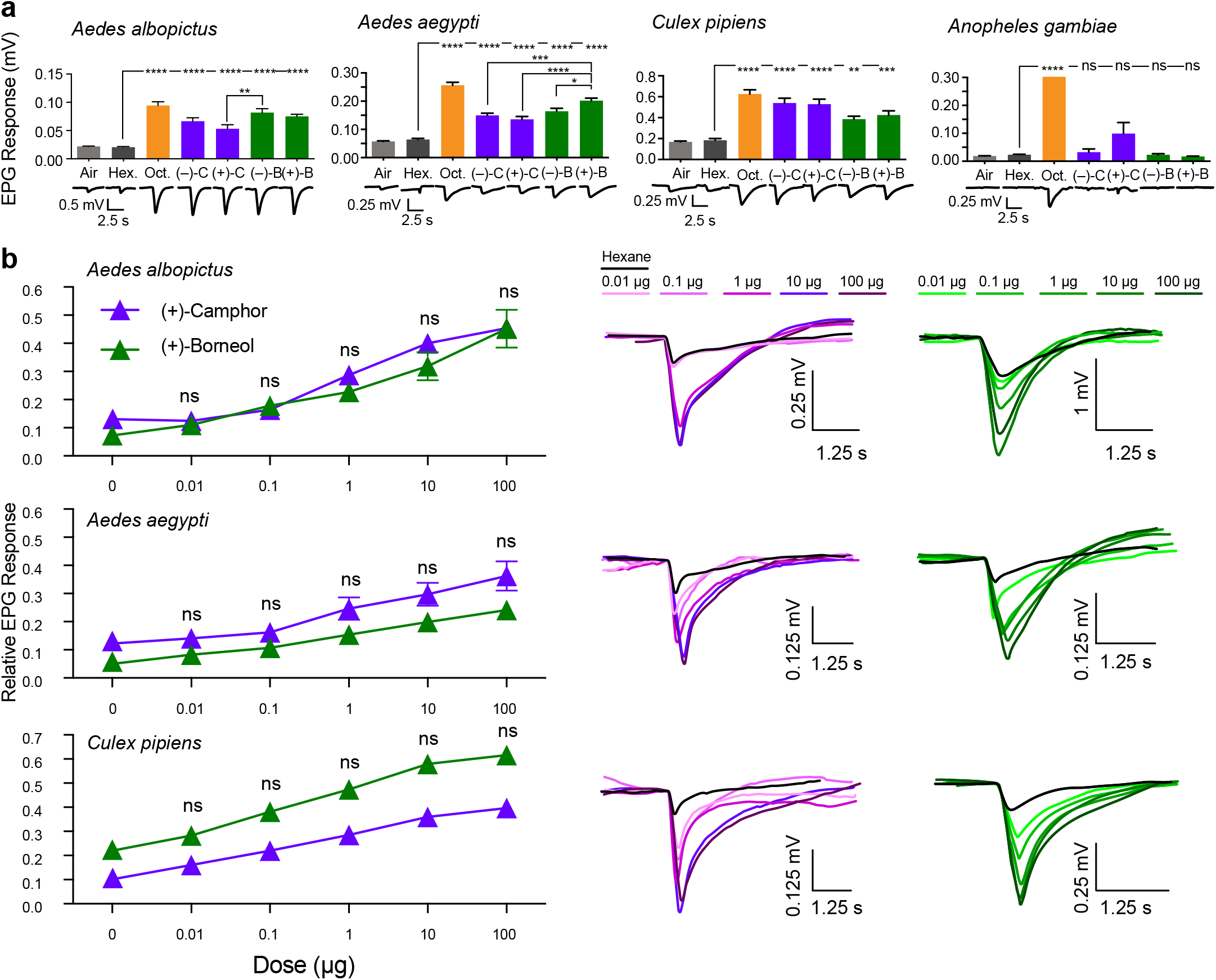
The maxillary palp of culicine mosquitoes respond to borneol and camphor. **a**, Electropalpogram (EPG) responses in *Aedes albopictus, Aedes aegypti, Culex pipiens*, and *Anopheles gambiae*. EPG responses to 1-octen-3-ol (blue, 10 μg), camphor (purple, 10 μg), and borneol (green, 10 μg) enantiomers relative to hexane. Representative traces for each odorant are shown below the x-axis. Statistical differences were evaluated via one-way anova. *p < 0.05, **p < 0.01, ***p < 0.005 and ****p < 0.001. Data indicate the means ± SEM, *n* = 15. **b**, Dose-response relationships of the maxillary palps of three culicine mosquito species in response to increasing concentrations of (+)-borneol and (+)-camphor. Representative traces are shown on the right. Statistical differences were evaluated by multiple t-tests. Data indicate the means ± SEM, *n* = 15.

To test a potential causal gene-function relationship between *Or49* and the borneol response, we used homology directed repair (HDR) to knock-in a transgenic construct containing the QF2 transactivator and a ubiquitously expressing eCFP fluorescent marker driven by the OpiE2 promoter at the start codon of the *Or49* gene in *Ae. aegypti* (Supplementary Fig. 6). To investigate the genetic mechanism determining borneol sensitivity in the palp, we conducted single-sensillum recordings (Fig. 4a) from the cp sensillum of *Ae. aegypti* (Fig. 4b), *An. gambiae* (Fig. 4c), *Ae. albopictus* (Fig. 4d), and *Cx. quinquefasciatus* (Fig. 4e). The response of the cpA and cpB neurons to CO_2_ ^22^ (data not shown) and *R*-(–)-1-octen-3-ol ^23^, respectively, has been well characterized in previous studies ^13–15^. (+)-Borneol elicited a dose-dependent response in the cpC neuron of *Ae. aegypti, Ae. albopictus, Cx. quinquefasciatus* but not in *An. gambiae s*.*s*. (Fig. 4f). Single sensillum recordings identified the spontaneous activity of three sensory neurons, distinguished by differences in spike amplitudes, housed in the capitate peg sensilla of wild type *Ae. aegypti* (Fig. 4c) and *An. gambiae* (Fig. 4d). No SSR responses were recorded from the cpC neuron in the *Ae. aegypti Or49* null mutant line in response to (+)-borneol (Fig. 4c) indicating that *Or49* is sufficient and necessary to confer the cpC neuron sensitivity to borneol. The lack of response of the cpC neuron in *Anopheles* (Fig. 4d and f) is consistent with the expression of *Or28* instead of *Or49*, which is not activated by borneol (Supplementary Fig. 5). The absence of OR28 activation and *An. gambiae* palp EPG responses by borneol and camphor support this conclusion.

**Figure 4.**
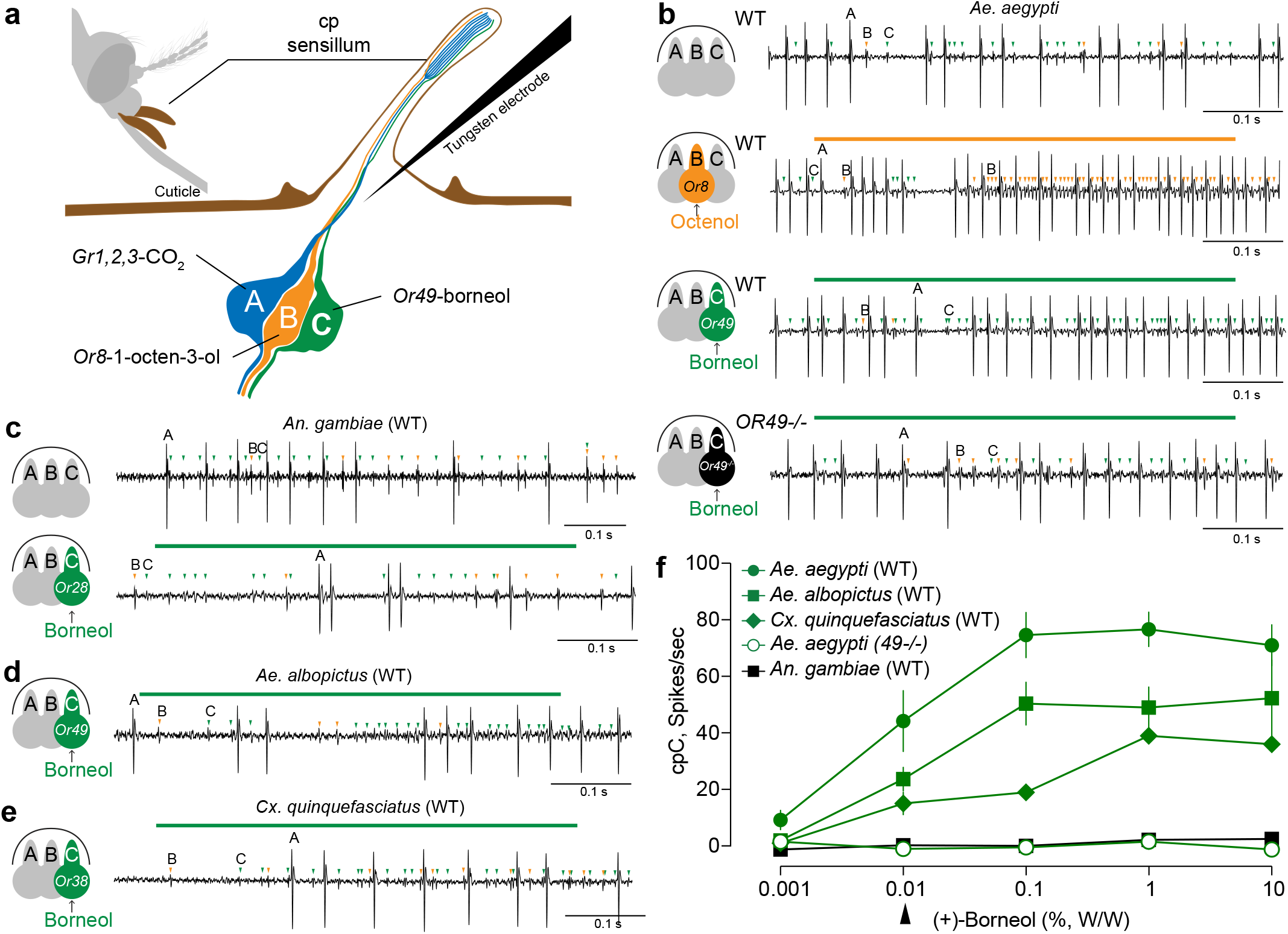
Species-dependent response to (+)-borneol of the C neuron of the capitate peg sensilla and its reliance of *Or49* in *Ae. aegypti*. **a**, A tungsten electrode was used to record action potentials (spikes) from the capitate peg sensillum on the maxillary palp, which houses three neurons. The cpA neuron responds to CO_2_ (large spike in traces), the cpB neuron responds to 1-octen-3-ol (medium size spikes) and the cpC neuron responds to borneol (small size spikes). **b-e**, The spontaneous activity of the three sensory neurons in the capitate peg sensilla of *Ae. aegypti* and *Anopheles gambiae s*.*s*.. In the top traces, note the differences in spike amplitude of the A, B (orange markers) and C neurons (green markers). *R*-(–)-1-octen-3-ol and (+)-borneol elicit responses in the B and C neurons of *Ae. aegypti*, respectively. The response to (+)-borneol is abolished in *AaegOr49*^*-/-*^ mutant mosquitoes. The C neuron of *An. gambiae s*.*s*. does not respond to (+)-borneol. **b-e**, Representative traces of the cp sensilla of *Ae. albopictus* and *Cx. quinquefasciatus*. **f**, Species- and dose-dependent response of the C neuron to (+)-borneol with mean ± s.e.m. at each concentration. The arrow and dotted line indicate the borneol dose to the shown representative traces. *Ae. aegypti* (*n* = 10), *Ae. albopictus* (*n* = 7), *Cx. quinquefasciatus* (*n* = 2), *Ae. aegypti Or49-/-* (*n* = 8), *An. gambiae* (*n* = 15).

### Borneol and camphor specifically activate the MD3 glomerulus in the antennal lobe

To examine how *Ae. aegypti* processes borneol in the antennal lobe (AL), two-photon imaging experiments were conducted using pan-neuronal GCaMP-expressing *Ae. aegypti* mosquitoes. We utilized existing mosquito lines that contained a *QUAS-GCaMP6s* transgene crossed with the *brp-QF2* driver line ^24^, allowing the directed expression of the calcium indicator GCaMP6s in all neurons of the AL. Mosquitoes were glued to holders that permitted two-photon imaging of calcium responses in the AL (Fig. 5a), allowing repeatable registration of the AL glomeruli between preparations (Fig. 5b) and recording of glomerular responses (Fig. 5c) ^25^. Glomeruli from our two-photon imaging results were mapped to AL atlases ^26,27^, allowing accurate identification of the glomeruli of interest in their spatial context (Extended Data Fig. 5). We focused on the mediodorsal (MD) glomeruli, as these glomeruli receive input from the capitate peg sensilla of maxillary palps (Fig. 5b and c) and are responsive to host odors, including 1-octen-3-ol (MD2) and CO_2_ (MD1) ^27^.

**Figure 5.**
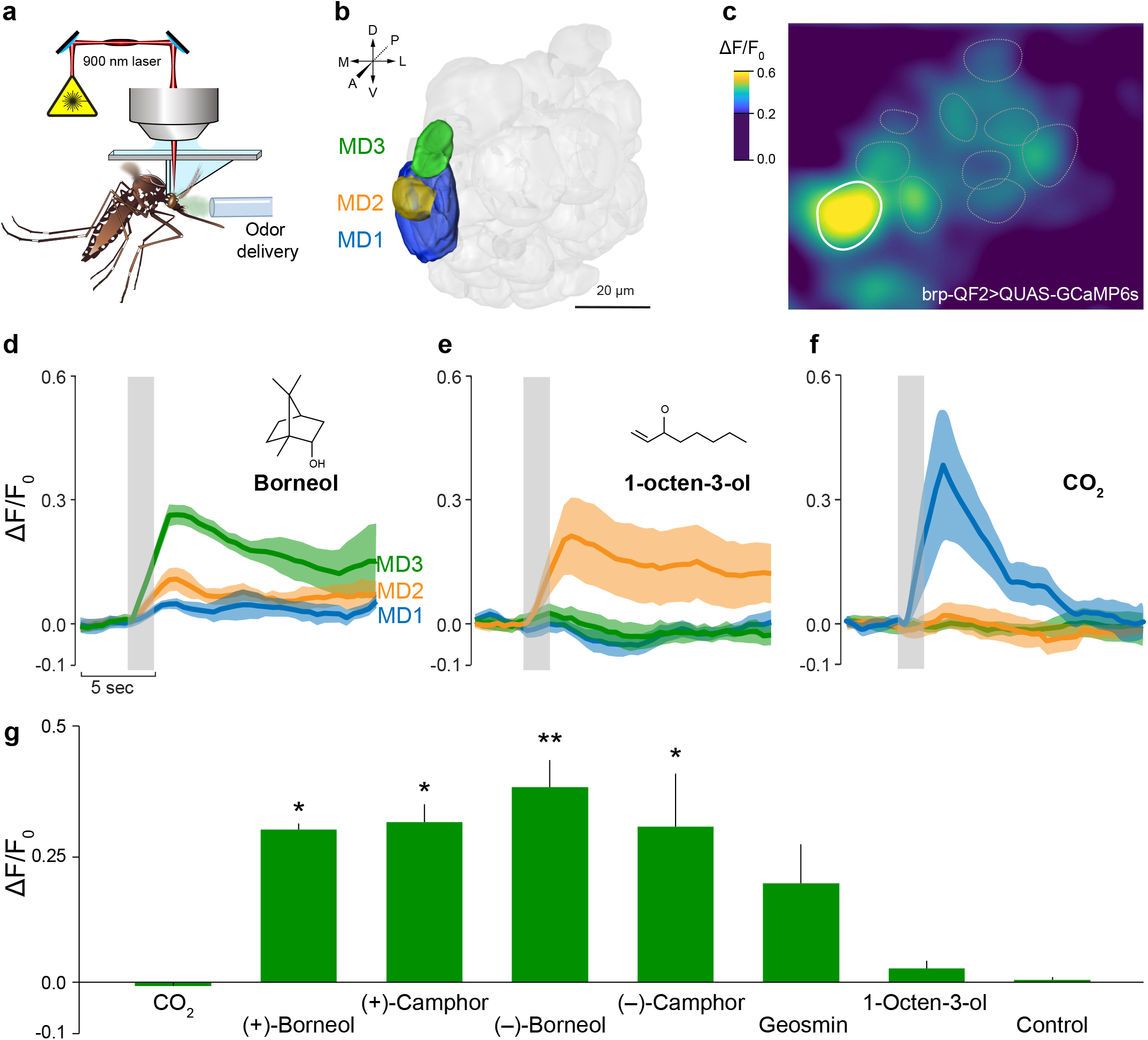
Borneol elicits robust responses in the *Ae. aegypti* AL. **a**, Schematic of the two-photon setup used to record calcium dynamics in the mosquito antennal lobe (AL). **b**, AL atlas, highlighting the MD1 (blue), MD2 (orange), and MD3 (green) glomeruli. Non-responsive AL glomeruli (grey) and the mediodorsal glomeruli were registered and mapped to previously published atlases. **c**, Pseudo color plot from a single preparation of ΔF/F_0_ calcium responses (0-0.8 scale) to (+)-borneol (10^−4^ dilution), at a depth of 75 µm from the surface of the AL. Borneol evoked a strong response in the region of interest mapped to the MD3 glomerulus (highlighted in white). **d**, Glomerular responses (ΔF/F_0_) to (+)-borneol for the MD1, MD2, and MD3 glomeruli. Lines are the mean of one glomerulus (*n* = 4-8 preparations); shaded areas are the SEM. The grey bar denotes stimulus duration (2 s). **e**, Same as in D, except the glomeruli were stimulated with 1-octen-3-ol (10^−4^ dilution). **f**, Same as in D, except the glomeruli were stimulated with CO_2_ (5%). **g**, Tuning curve for the MD3 glomerulus to a limited panel of 8 odorants, each tested at 10^−4^ concentration. The MD3 glomerulus (bars in green) showed significant calcium responses to enantiomers of borneol and camphor compared to the solvent control (Kruskal-Wallis test: χ^2^ = 37.1, P < 0.0001; posthoc multiple comparisons: p < 0.05). Bars represent the mean ± SEM.

To determine the odor coding of the MD glomeruli, and identify the cognate glomerulus representing borneol, we first recorded from the MD1-3 glomeruli while stimulating with CO_2_ (5%), 1-octen-3-ol (10^−4^ dilution), borneol (10^−4^ dilution), and the solvent control. For the MD3 glomerulus, (+)-borneol elicited strong, tonic responses that were significantly greater than the solvent control (P<0.001) (Fig. 5d and g). By contrast, responses of the MD1 and MD2 glomeruli to borneol were not significantly different from the control (P=0.09 and 0.23, respectively). The MD2 glomerulus showed the greatest responses to 1-octen-3-ol (Fig. 5e) (P = 0.00003), and the MD1 glomerulus to CO_2_ (P=0.01) (Fig. 5f). Given MD3’s response to (+)-borneol, and OR49/cpC’s responses to other terpene compounds, we next examined how this glomerulus responded to a limited panel of different odorants, including enantiomers of borneol and camphor. From this panel, (–)-borneol elicited the greatest response in the MD3 glomerulus (Fig. 5g), closely followed by (+)-borneol, enantiomers of camphor, and geosmin. These findings are consistent with previous studies demonstrating that the three MD glomeruli receive input from the maxillary palp cp neurons whereby the MD1 glomerulus receiving CO_2_ input from the cpA neurons ^28^ while the medium size MD2 glomerulus receives 1-octen-3-ol input from the cpB neurons ^27^.

### Borneol repels human-host seeking female *Ae. aegypti*

To examine the impact of (±)-borneol on blood-seeking *Ae. aegypti* mosquitoes, we conducted an arm-in-a-cage assay (Extemded Data Fig. 1a), exposing 15 females to human skin odor for 10 minutes (Figure 6). A protective glove covered the hand, allowing mosquitoes to detect the odor through a dorsal open area, which was protected with a screen and equipped with a chemical holder for the deposition of (±)-borneol (1M) (Extended data Fig. 1). To monitor host-seeking behavior—specifically, the presence of females, their walking path (in cm), and visit duration in the region of interest (ROI)—we fine-tuned a custom object detection YOLOv8 model ^29^. Our results demonstrated a significant reduction (54%) in the number of trajectories within the ROI when the hand was treated with (±)-borneol compared to the vehicle-treated hand (Fig. 6a). All trajectories across treatments and repetitions are shown in Figure 6b, where different colors indicate different trajectories. Vehicle and borneol-induced trajectories were 951 and 428, respectively This trend was consistent across time (Fig. 6c). On average, the total time spent in the ROI was three times higher in the control than in the (±)-borneol treatment (Fig. 6d). Additionally, the distance walked within the ROI was significantly lower in the presence of (±)-borneol (Fig. 6e). Figure 6f visually represents the mosquito detections throughout the experiment, highlighting the differences between the control and (±)-borneol treatments. A video sample is available in Extended Data Fig. 6. When purifying the *Or49* homozygote mutant, we noticed severe reduction in fitness, which precluded further behavioral experiments.

**Figure 6.**
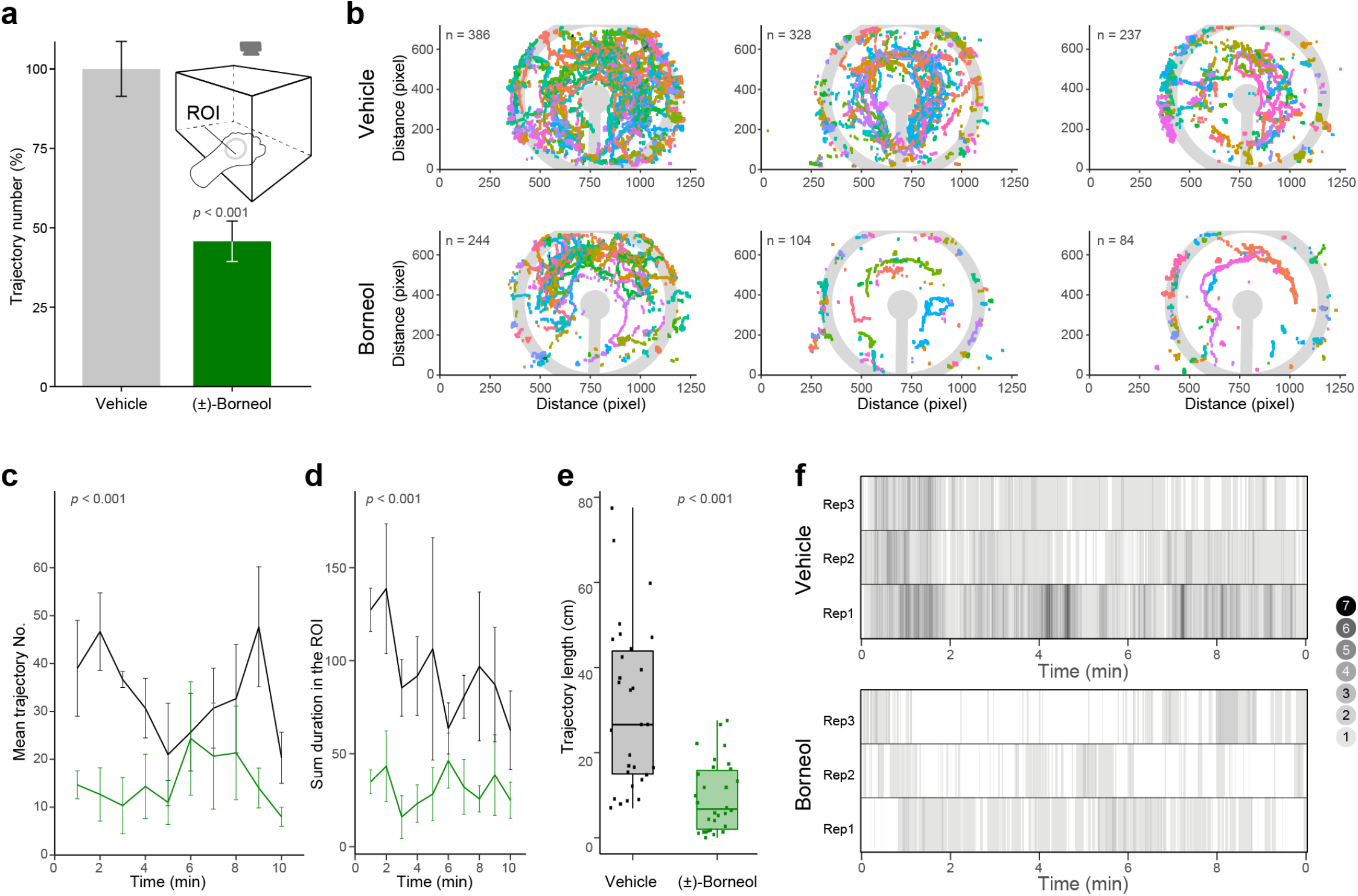
Borneol inhibits host-seeking female *Ae. aegypti* (Liverpool) behavior. **a**, The average number of females trajectoreis detection per minute (normalized to the control group) in the ROI in response to control (DEE) and treatment (DEE + racemic borneol). **b**, Schematics of all trajectories across treatment and repetitions (n=3). **c**, Average trajectories over time (min). **d**, Sum of trajectory durations (seconds) within the ROI over time (minutes). **e**, Total distance traveled per minute in the ROI. All comparisons were conducted with 30 data points for each treatment and tested with a Mann–Whitney U test (p-value < 0.001, *n* = 3). **f**, Schematics of all detections as a function of time.

## Discussion

We have identified a conserved mechanism in culicine mosquitoes responsible for the selective detection of borneol, a bicyclic monoterpenoid that has been used as an insect repellent since ancient times. Recent studies have shown that borneol is a broad-acting insect repellent against fungus gnats ^30^, the booklouse *Liposcelis bostrychophila* ^31^ and fire ants ^32^ suggesting that more than one olfactory-mediated mechanisms in insects are at play since *Or49* homologs in these insects have not been identified. Interestingly, borneol elicits responses in the *Ae. aegypti* antenna as well ^33^. However, antennal activation by borneol requires doses 10,000 times greater than in the maxillary palp (Supplementary Fig. 7). This difference in sensitivity in favor of the palp indicates that whatever mechanism is involved in the antenna is likely not an ecologically-relevant response but rather the product of chemical overstimulation. Moreover, this alternative mechanism is *Or49*-independent since its expression has not been reported in the antenna.

In *Anopheles*, the antennal-expressed *Or48* ^34^, one of the two closest homologs to the culicine *Or49* (Supplementary Fig. 1c), responds to straight chains alcohols, ketones, and acetates ^18,35^ suggesting that this receptor is tuned to a different class of compounds setting apart the olfactory coding logic between these two mosquito subfamilies. Contrary to OR8, which discriminates between the 1-octen-3-ol enantiomers ^21^, the two borneol enantiomers elicited comparable activations at the pharmacological, physiological and AL levels. These findings suggest that the receptor binding pocket accommodates both forms of borneol or that a closely related structural analog of borneol, with greater potency, remains to be identified^36^.

Our results suggest that in *Ae. aegypti* and other culicine mosquitoes, the repellent properties of traditional Chinese medicinal plants such as the camphor tree EO are largely mediated by the activation of OR49 by bicyclic monoterpenoids and more specifically by borneol. Borneol repels human host-seeking *Ae. aegypti, Cx. quinquefasciatus* and *An. stepehensi* ^33,37,38^ while its presence in EOs attracts gravid *Ae. aegypti* ^39^, suggesting that this compound has opposite effects on animal-host-seeking behavior and oviposition preferences. The reason for borneol being a signal mediating repellency is puzzling. While several monoterpenoids such as 1-8-cineol, limonene, and fenchone exhibit anticholinesterase activity ^40,41^ causing paralysis and death in a variety of insects, there are conflicting reports on the larvicidal activity of borneol and camphor against mosquito larvae ^39,42–45^. Future studies should focus on the ecological role of borneol in different contexts, including foraging and oviposition site selection.

We found that the co-location of the three cp neurons in the cp sensillum is reflected by the anatomical proximity of their projections in the MD glomeruli cluster in the AL. Whether the grouping of these three glomeruli has a functional significance and why the cp sensillum houses ORNs tuned to animal and plant host olfactory cues should be the focus of future studies. The two largest cp neurons are tuned to animal-host attractants whereas the smallest and third cpC neuron responds to a plant-host odorant with repellent activity. This antagonism in valence whereby an attractive signal is detected by the largest of two ORNs within the same sensillum while a deterring signal is detected by the smallest ORN is widespread in animals^46^. We surmise that the main function of the mosquito cp sensillum is to detect animal host odorants, while the cpC neuron exerts a presynaptic inhibitory effect mediated by lateral coupling when activated by borneol.

Our findings indicate that the cp sensillum in the palp of culicine mosquitoes detects signals from animal and plant hosts. Why OR49 is selectively tuned to this particular compound and not to a more ubiquitous volatile phytochemical remains a mystery. Further experiments will reveal how these three signals are integrated at the pre-synaptic level within the sensillum and in the AL. Understanding the ecological and neurological significance of the mosquito cp sensillum, i.e., the packaging of animal and plant host detectors within the same sensillum, may be exploited to design novel repellent formulations.

## Methods

### Insects

Our mosquito colonies were reared according to previously published protocols ^47^. *Ae. aegypti* originated from a colony established by Prof. Joel Margalit. The *Ae. aegypti* Liverpool strain was provided by the Akbari lab. *Aedes albopictus* was the FPA Foshan strain collected (Foshan, China) reared at the insectary of the University of Pavia since 2013 (Palatini et al., 2017). *Anopheles gambiae* was the G3 strain originally isolated from West Africa (MacCarthy Island, The Gambia) in 1975 (Federica Bernardini et al., 2017). *Culex pipiens* originated from a wild type population was provided by Dr. Laor Orshan (Ministry of Health, Israel).

### Phylogenetic and genomic analyses

The AaegOR49 protein sequence (AAEL001303-PA) was used as a query to identify homologs in other mosquito species. Amino acid sequences of homologous odorant receptors were obtained from the Vectorbase database (vectorbase.org) and the *Tx. Amboinensis* genome assembly (Zhou et al., 2014). The multiple sequence alignment was conducted using ClustralW (Chenna et al., 2003). The Maximum-likelihood phylogenetic tree was constructed using MEGAX software (Model: G, bootstraps: 5000). Exon-intron structure analysis was conducted manually by aligning predicted amino-acid sequences to genomic regions. Approximate mapping of the genes in chromosome location for each mosquito species and construction of synteny maps done using Geneious prime software (Kearse et al., 2012) using latest genome assemblies available for each mosquito species (Matthews et al., 2018, Palatini et al., 2020, Boyle et al., 2021, Arensburger et al., 2012, Sharakhova et al., 2007).

Genomic DNA (gDNA) was isolated from whole bodies of *Tx. amboinensis* adults. gDNA was diluted to 25ng/µL in nuclease free water and used as a template in polymerase chain reactions to amplify odorant receptors (TambOR) using Taq polymerase and the following primer sets: TambOr6.F1 (ATGCGCTTCTACGAGAAATAC), TambOr6.R1 (TCAGAAATTATCCTTCAGGATC); TambOr12.F1 (ATGCCATCGGTTTTCTTGGTT), TambOr12.R1 (CTAAAACACTCGCTTCAATATC); TambOr13.F1 (ATGTTCTGCTTCAGGAAGATC), TambOr13.R1 (CTAGAAGTGGTTTTTCAATATAA); TambOr49.F1 (ATGTTGTTCAAGAACTGTTTCC), TambOr49.R1 (TTAATAATTGAATCTTTCCTTCAG); TambOr71.F1 (ATGGGCAGCAGTGATGGTGAC), TambOr71.R1 (CTACTGGTTGATTTTACTGAGG).

Cycling conditions were 94°C for 60s; 30 cycles of 94°C for 20s, 56°C for 20s, 72°C for 30s; and a final extension of 72°C for 5 minutes. Amplicons were analyzed by electrophoresis on a 1% agarose gel, cloned into the TOPO-TA pCR2 plasmid, heat-shock transformed into TOP10 competent *E. coli* cells, and grown overnight at 37°C on LB+ampicillin+X-gal agar plates. Ampicillin resistant, white colonies were isolated using a sterile pipette tip and grown overnight 37°C in 3mL of LB+ampicillin. Plasmids were isolated by DNA miniprep and the Sanger method was used to determine the DNA sequence in both directions using standard T7 and M13.rev primers. TambOR nucleotide sequences were compiled into contigs and intron/exon regions were inferred by comparing to coding sequences described in a previous publication ^48^.

### Chemical reagents

The chemicals used for the deorphanization of receptors were obtained from Acros Organics (Morris, NJ, USA), Alfa Aesar (Ward Hill, MA, USA), ChemSpace (Monmouth Junction, NJ, USA), Sigma Aldrich (St. Louis, MO, USA), TCI America (Portland, OR, USA), and Thermo Fisher Scientific (Waltham, MA, USA) and Penta Manufacturing Corp. (Livingston, NJ, USA) at the highest purity available. *Cannabis* essential oils and sub-mixtures were formulated and supplied by Eybna Technologies (Givat Hen, Israel) (Table S1 and Supplementary Fig. 2–3).

### Two-electrode voltage clamp of *Xenopus laevis* oocytes

In vitro transcription and two-microelectrode voltage clamp electrophysiological recordings were carried out as previously described ^49^. Experimental procedures for the *Ae. albopictus* receptor clone was performed as previously described ^50^. *AalbOr49* and *AalbOrco* templates were synthesized by Twist Biosciences (San Francisco, CA, USA) and cloned into the pENTR^*TM*^ vector using the Gateway^*R*^ directional cloning system (Invitrogen Corp., Carlsbad, CA, USA) and subcloned into the *X. laevis* expression destination vector pSP64t-RFA. Several of these genes were codon-optimized (Supplementary Table 2). For pairwise current comparisons of (+)-borneol and (+)-camphor, two separate perfusion systems were assembled, one for each compound. The outlet from each system was connected to a 2 to 1 perfusion manifold. At each tested concentration, (+)-borneol and (+)-camphor were administered consecutively. Representative current traces can be seen in Supplementary Figure 4. *Odorant receptor 49* and *Orco* clones and recording measurements can be found in Supplementary Table 2 and 3, respectively. The use of *Xenopus laevis* frog eggs was carried out according to The Hebrew University of Jerusalem Ethics committee.

### Electropalpogram recordings

Based on the voltage clamp data, responses from the maxillary palps of *Ae. albopictus, Ae. aegypti, Cx. pipiens* and *An. gambiae* were recorded with camphor or borneol. Electopalpogram (EPG) assays were performed according to published procedures ^51^ with a few modifications. Within the Pasteur pipette, 20 μL of odorants diluted in hexane were deposited at desired dose onto a 4×20 mm Whatman filter paper strip. The odorants were delivered into a consistent humidified airstream (at a flow rate of 50 cm/s) at approximately 2 cm from the maxillary palps. The pulse duration was 0.1 s, and the recording time was set for 5 s. A 2-min gap was allowed between stimuli to recover the EPG sensitivity. At least 5 individual females were tested, for each individual, 3 technical replicates were performed with doses ranging from 0.01 ug to 100 μg. Negative controls (hexane and air) and positive controls (1-octen-3-ol) were performed at both the beginning and end for each recording session to monitor the decline in sensitivity of the maxillary palps. EPG peak responses were normalized to the response of hexane.

### Generating *Or49* knockout line in *Ae. aegypti* and isolating an *OR49*^*-/-*^ homozygous line

To knock out the gene encoding the *Ae. aegypti* OR49 receptor, we used homology directed repair (HDR) to knock in a transgenic construct containing the QF2 transactivator and a ubiquitously expressing eCFP fluorescent marker driven by the OpiE2 promoter at the start codon of the *Or49* gene (Supplementary Fig. 6a–b). HDR was triggered by double strand breaks at the *Or49* promoter and the 1^st^ exon mediated by two guide RNAs, whose cleavage activities had been tested *in vitro* (Supplementary Fig. 6c). Genomic DNA was extracted from whole bodies of *Ae. aegypti* Liverpool individuals using the DNeasy Blood & Tissue Kit (Qiagen, Redwood City, CA). Homology arms of the *Or49* gene flanking 1 kb upstream and downstream of the desired insertion site (Supplementary Fig. 6a) were amplified with the Q5 High Fidelity DNA polymerase (New England Biolabs, Ipswich, MA), using primer pairs Left_Arm_Or49_FWD and Left_Arm_Or49_REV as well as Right_Arm_Or49_FWD and Right_Arm_Or49_REV, respectively (Supplementary Table 4). The QF2-ECFP DNA cassette was amplified using the V1117A plasmid as template and the primer pair QF2_OpIE_ECFP FWD and QF2_OpIE_ECFP REV. The cassette contains the *QF2* sequence and the 3’ UTR of the *HSP70* gene along with the *ECFP* gene under the control of the *Opie2* promoter and the 3’ UTR of the *SV40* gene. PCR bands were cut out from agarose gels and purified with Zymoclean Gel DNA Recovery Kit (Zymo Research, Irvine, CA).

Upstream and downstream homology arms as well as the QF2-ECFP cassette PCR products were assembled into the backbone of the V1117A plasmid (Supplementary Fig. 6b) using Gibson assembly reaction, following the manufacturer recommendations. Gibson’s reaction-derived plasmid product was used to transform JM109 cells (Zymo Research), and colonies were individually grown overnight for minipreps with Zyppy Plasmid Miniprep Kit (Zymo Research). The sequence of the plasmid V1117F-Or49 was confirmed by restriction enzyme digestion using the enzymes ScaI-HF and AvrII as well as by whole plasmid sequencing (Primordium Labs, Monrovia, CA). V1117F-Or49 plasmid was used to retransform JM109 cells, which were grown for maxiprep purification using the PureLink Expi Endotoxin-Free Maxi Plasmid Purification Kit (Thermo Fisher Scientifics, Waltham, MA).

For guide RNA synthesis, non-template reactions were carried out with the gRNA_Left_Or49_F and gRNA_Left_Or49_R forward primers and the universal guide RNA reverse primer (Universal-sgRNA_R). PCR bands were isolated from agarose gel and purified as above described. Guide RNAs were synthesized with the Ambion MEGAscript kit (Thermo Fisher Scientifics) for 4 hours at 37°C, using 300 ng of purified PCR product. Guide RNAs were further purified with the Megaclear Kit (Thermo Fisher Scientifics).

To assess the cleavage activity of the synthesized guide RNAs, *in vitro* Cas9 cleavage assays were carried out. A DNA fragment spanning 1,009 bp overlapping the cleavage sites of the guide RNAs was amplified with primers Or49_cleav_F and Or49_cleav_R, DNA bands were isolated from agarose gel, purified, and 100 ng of which was used in cleavage assays with 300 ng of recombinant Cas9 (PNA BIO, Thousand Oaks, CA) and 100 ng of each guide RNA upon incubation at 37°C for 1 hour (Supplementary Fig. 6c).

For embryo microinjection, an injection mix was prepared with the plasmid V1117F-Or49 at 500 ng/ul, gRNAs left and right at 100 ng/ul each, and recombinant Cas9 at 300 ng/ul. Mix was filtered with the Ultrafree-MC Centrifugal Filter UFC30GUOS (Millipore, Burlington, MA). Freshly harvested *Ae. aegypti* embryos (Liverpool strain) were injected with the transformation mix using a custom-made insect embryo microinjector (Hive Technologies, Cary, NC) at 100 psi pulse and 3 psi back pressures, using quartz needles pulled with a P-2000 needle puller (Sutter Instrument Co., Novato, CA). Genomic DNA of the G1 fluorescent mosquitoes (Supplementary Fig. 6d) were individually extracted with the Qiagen blood kit, and PCR amplified for Sanger Sequencing reactions (Retrogen, San Diego, CA) using the primer pairs Or49_diag_up_F and Or49_diag_up_R targeting the upstream and primer pair Or49_diag_down_F and Or49_diag_down_R targeting the downstream insertion site (Supplementary Fig. 6e). For whole DNA cassette sequencing (Primordium), 2.5kb DNA fragments were amplified from the DNA of G1 individuals using the primer pair Or49_diag_up_F and Or49_diag_down_R (Supplementary Fig. 6f).

For isolation of homozygous knockout individuals, pupae were sex sorted, and single pairs were transferred into Narrow *Drosophila* Vials (Genesee Scientific, El Cajon, CA), filled with 10 mL of deionized water (DI) and closed with cotton stoppers. Upon emergence, water was drained, and sugar cottons were placed into the vials. Mosquitoes were allowed to mate for 5 days in the vials, and then all individuals were transferred to a single cage for blood feeding. Three days after blood feeding, females were individually transferred to a fresh vial containing a piece of brown paper towel wetted with 1 mL DI water. Females were allowed to lay eggs, and each egg batch was individually hatched in 250 mL Clear Pet Cups (9oz). Larvae were screened for fluorescence, and the batches that resulted in 100% fluorescent individuals were grown to adulthood and intercrossed to confirm homozygosity in the following generation. Two out of thirty egg batches resulted in homozygous offspring. To further confirm the homozygous status of these individuals, genomic DNA of three pools of 10 males, 10 females, and a mix of males and females were PCR amplified with the primer pair Or49_diag_up_F and Or49_diag_down_R, which resulted in a single band for homozygous individuals and two bands for heterozygous individuals (Supplementary Fig. 6g). Out of 739 injected embryos, 5.5% survived to the pupal stage. These G_0_ individuals were sex sorted and outcrossed with wildtypes. Sixteen G_1_ larvae displayed blue (cyan) fluorescent bodies (Supplementary Fig. 6d). Insertion of the transgenes within *Or49* was confirmed in multiple G_1_ individuals (Supplementary Fig. 6e–f) by sequencing of the insertion sites as well as the transgene (Supplementary Fig. 6e–f) resulting in a nonfunctional receptor. An *Or49* ^-/-^ homozygous mosquito strain was isolated by single-pair mating, screening the fluorescence marker in the offspring for 100% penetrance, and PCR confirmation for the presence of homozygous mutant alleles (Supplementary Fig. 6g).

### Single-sensillum electrophysiology

Single sensillum recordings from capitate peg sensilla on the maxillary palp of wild type *Ae. aegypti* (Liverpool), *Ae. albopictus* (FPA), *Cx. quinquefasciatus* (Thai) and *An. gambiae sensu stricto* (G3) and *Ae. aegypti Or49*^*-/-*^ mutant mosquitoes were performed using an established protocol ^52^. Spikes were quantified offline using the established nomenclature for the sensory neurons ^53^. The number of spikes counted during a 0.5 s stimulus delivery interval was subtracted from the number of spikes counted during a 0.5 s prestimulus period, and the result was multiplied by 2 to obtain the activity of individual sensory neurons housed in the capitate peg sensillum as a spikes/s measurement.

To investigate the physiological activity of the A, B and C neurons housed in single capitate peg sensilla, CO_2_, *R*-(–)-1-octen-3-ol and (+)-borneol were used: gas cylinders containing metered amounts of CO_2_ (300, 600, 1200, 2400, or 4800 ppm) and oxygen (20%), balanced by nitrogen (Strandmöllen AB, Ljungby, Sweden) were used to assess the activity of the A neuron; serial decadic dilutions of *R*-(-)-1-octen-3-ol (CAS: 3687-48-7, Penta Manufacturing, Livingston, USA), diluted in paraffin oil, were used to assess the activity of the B neuron; and serial decadic dilutions of (+)-borneol (CAS: 464-43-7, Sigma-Aldrich, St. Louis, MO, USA), diluted in diethyl ether (SupraSolv, Billerica, MA, USA) were used to assess the activity of the C neuron. Pasteur pipettes were filled with the metered CO_2_ and used immediately to stimulate the preparation. A 15 μL aliquot of each dilution of *R*-(–)-1-octen-3-ol and (+)-borneol was pipetted onto a filter paper (5 mm × 15 mm) inserted inside a Pasteur pipette, and the diethyl ether was allowed 15 min to evaporate, before being used for stimulus delivery. All stimuli were delivered into the airstream passing over the maxillary palp preparation.

### Calcium imaging in the *Ae. aegypti* antennal lobe (AL)

Odor-evoked responses in the *Ae. aegypti* antennal lobe (AL) were imaged using the brp-QF2>QUAS-GCaMP6s progeny from the brp-QF2 and QUAS-GCaMP6s parental lines ^24^. A total of eight 6-8 day-old female mosquitoes were used for all calcium experiments. Each mosquito was cooled on ice and transferred to a Peltier-cooled holder that allowed the mosquito head to be fixed to a custom stage using ultraviolet glue. The stage permits the superfusion of saline to the head capsule and space for wing and proboscis movement ^25,54^. Once the mosquito was fixed to the stage, a window in its head was cut to expose the brain, and the brain was continuously superfused with physiological saline ^55^. Calcium-evoked responses in the AL were imaged using the Prairie Ultima IV two-photon excitation microscope (Prairie Technologies) and Ti-Sapphire laser (Chameleon Ultra; Coherent; at 1910 mW power). Experiments were performed at 75 µm depth from the ventral surface of the AL, allowing characterization of the mediodorsal glomerular responses to olfactory stimuli and allowing these glomeruli to be repeatedly imaged across preparations. To record odor-evoked responses, images were collected from a 110 µm × 83 µm plane at 2 Hz (line period of 1 ms), and for each odor stimulus, images were acquired for 25 s, starting 10 s before the stimulus onset. Image data were imported into Matlab (v2017; Mathworks, Natick, Massachusetts) for Gaussian filtering (2×2 pixel; σ = 1.5-3) and alignment using a single frame as the reference at a given imaging depth and subsequently registered to every frame to within ¼ pixel. Odor stimuli were diluted to 10^−3^ and 10^−4^ concentration in hexane (>99.5% purity; Sigma), with hexane used as the solvent control. During an experiment, odor stimuli were separated by intervals of 120 s to avoid receptor adaptation, and odor syringes were used once per preparation to prevent decreased concentration within the cartridge. Calcium responses are calculated as the change in fluorescence and time-stamped and synced with the stimulus pulses. After an experiment, the AL was sequentially scanned at 0.5 µm depths from the ventral to the dorsal surface to provide glomerular assignment and registration between preparations. Glomeruli (1 µm^3^ voxel) were mapped and registered based on the positions and odor-evoked responses of the putative AL3, MD1-3, and AM2 glomeruli, using available AL atlases ^26,27^ and the Amira software (v. 6.5, FEI Houston Inc.).

### Behavioral assay

The role of borneol in human host-seeking female mosquitoes was examined with an arm-in-a-cage assay (Extended Data Fig. 1a) described previously ^47^ with minor modifications (Extended Data Fig. 1b and c) in an air-conditioned room (26±1 °C, 60±5 % RH). Briefly, the experimenter’s hand was presented to fifteen 5–10 days post emergence adult females. A three-dimensional-printed interlocking ring with a diameter of 55 mm was used in this experiment (Extended Data Fig. 1b). The ring was placed over the dorsal side of a nitrile glove (powder-free latex). To evoke human odor and prevent mosquito bites, we replaced the nitrile glove between the two ring components with a plastic net (Extemded Data Fig. 1c). The interlocking ring included a central odorant delivery platform comprising a 10-mm-diameter cover glass and two 5-mm-diameter filter discs (WHA10016508; Merck) for the evaluation of VOCs (stl files are provided in Extended Data Fig. 2 and 3). Plastic net, nitrile glove and odorant delivery platform were replaced between repetitions to avoid contaminations. Mosquitoes were placed in a 20.3-cm^3^ metal cage located in an experimental room with a vent, under a video camera (EOS 70D, lens: MACRO 0.25/0.8ft; Canon Inc., Tokyo, Japan) and a light ring. Mosquitoes were allowed to acclimate for ten minutes before recording. Mosquito behavior was recorded for 10 min at 25 frames per second. The number of mosquito detection on the screen and ring were automatically counted using a custom YOLOv8 model. We used the solvent diethyl ether (DEE) as a vehicle. On a blank filter disc, 25 μL of DEE, and DEE with racemic borneol (1M, 3.58 mg) were deposited on the filter paper and allowed to evaporate for 2 min prior to mosquito exposure outside the experimental room to avoid contamination. To enhance attractiveness, at the beginning of each experiment, the experimenter first rubbed the ring-mounted glove against the shirt and skin for 1 minute. Additionally, the experimenter blew twice, once into the cage and once into the glove. All experiments were conducted during the first 4 hours of the diurnal period and lasted 10 min. This schedule was chosen for practical reasons and because mosquitoes consistently exhibited attraction to the human hand. Mosquito detection was normalized to the mean detection of the control 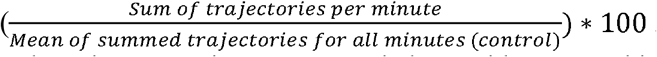. Statistical comparisons of mosquito detections per minute were carried out with Mann Whitney U test (*P*-value < 0.001, *n* = 3). For data variation across time and repetitions, see Figure 6. Data was analyzed using R Studio ^56^.

### YOLOv8 model

In this study, we fine-tuned the pre-trained YOLOv8m model ^57^using a customized dataset comprising images from the bioassay, along with additional data from previously published literature ^58^, to investigate the behavioral role of borneol on mosquitoes. The training process involved 250 epochs with a batch size of 10, utilizing separate sets of images and corresponding annotations for training, validation, and testing (5,334, 1,498, and 1,479 images, respectively). To assess the model’s performance, we employed the mean Average Precision (mAP) metric across a range of intersection-over-union (IoU) thresholds from 50% to 95% (mAP50-95). The resulting mAP50-95 value of 0.755 indicates the average precision considering different IoU thresholds. Furthermore, the recall value of 0.978 indicates a high proportion of true positive detections in this setup, while the precision value of 0.978 reflects the accurate identification of mosquitoes. It’s worth noting that detections were collected with a minimum confidence threshold and IoU of 0.5 to ensure reliable results. Video samples with automated detection are available in Extended Data Fig. 6. The model output was proccesed with Python, resulting dataframe with the following columns, including frame, x, y, treatment and trajectory identification. Statistical comparisons of all parameters were carried out with Mann Whitney U test with Benjamini-Hochberg (BH) adjustment method. Data was analyzed using R Studio (Team, 2021). The behavioral data can be found in (Extended Data Fig. 4). All codes can be obtained by request to the corresponding authors.

## Supplementary information

**Supplementary Figure 1. The phylogenetically-conserved gene encoding OR49. a**, Amino-acid sequence alignment of OR49 homologs in *Culex quinquefasciatus* (Cqui), *Aedes albopictus* (Aalb), *Aedes aegypti* (Aaeg), *Toxorhynchites amboinensis* (Tamb), and *Anopheles gambiae* (Agam). The top chart presents the amino-acid consensus. High consensus marked in green to yellow, low consensus are in red. **b**, Amino-acid sequence identity matrix. Intensity of shading indicates percentage of homologies. **c**, Phylogenetic tree of mosquito OR49 proteins (green). Closest brachyceran homologs (red branches) include *Glossina fuscipes* (Gfuc) and *Drosophila melanogaster* (Dmel). Bootstrap support above 50% are shown. Gene exon structure of OR49 genes and other phylogenetically-related genes. **d**, Exon compoistion of the OR49 group and related homologs. Exons are labeled in white and black. The nucleotide positions of exon-exon boundaries are shown and intron phases are color-coded. **e**, Chromosomic locations of Or49 homolog genes (green) and indolergic ORs (indolORs, in red). **f**, Syntenic relationships of OR49 genes in mosquito species and gene structure of OR49 genes.

**Supplementary Figure 2. List of volatile organic compounds present in *Cannabis* essential oils**. Each Cannabis blend represented by a distinct color is composed of VOCs belonging mainly to different terpene categories but also to non-terpene chemical classes.

**Supplementary Figure 3. Representative current traces of oocytes expressing mosquito OR49 proteins. a**, Representative traces of TambOR49+TambOrco injected oocytes vs monoterpenoid blends and compounds. Pineapple Haze (PH), Adom #9 (A9), O.G Kush (OGK), Master Kush (MK) and Jack Herrer (JH) triggered a measurable response. (±)-camphor, (+)-camphor, (+)-fenchone and eucalyptol were used as positive controls at concentration of 0.1 mM, all *Cannabis* essential oils were diluted by a factor of 1.5×10^5^. **b**, Representative trace of TambOR49+TambOrco vs sub-mixtures No. 1 through 6, which were formulated based on overlapping active compounds from *Cannabis* essential oils. All sub-mixtures were diluted by a factor of 1.5×10^6^. Only sub-mixture No.2 activated TambOR49+TambOrco. (+)-Camphor was used as positive control at concentration of 0.1 mM. **c**, Representative trace of TambOR49+TambOrco vs the components of sub-mixture No. 2. (±)-Borneol elicited currents 8-fold larger than the other tested single monoterpenoids. All compounds were tested at concentration of 0.1 mM. **d**, Representative traces of TambOR49+TambOrco concentration response curves (CRCs). Top left TambOR49+TambOrco vs (-)-borneol, Top right TambOR49+TambOrco vs (+)-borneol and bottom left TambOR49+TambOrco vs (+)-camphor. **e**, Representative traces of AaegOR49 CRC’s. Left AaegOR49 vs (–)-Borneol; right, AaegOR49 vs (+)-Borneol. **f**, Representative traces of CquiOR38+CquiOrco CRCs in response to increasing concentrations of (+)-camphor (left) and (+)-borneol (right). **g**, Representative traces of AaegOR49+Orco CRCs in response to increasing concentrations of (+)-camphor (left) and (+)-borneol (right). **h**, Representative traces of AalbOR49+AalbOrco CRCs in response to increasing concentrations of (+)-camphor (left) and (+)-borneol (right). **i**) Representative traces of pairwise current comparisons between (+)-camphor and (+)-borneol. From top to bottom, TambOR49+TambOrco, CquiOR38+CquiOrco, AaegOR49+AaegOrco and AalbOR49+AalbOrco. Down arrows indicate odorant administrations and concentrations. All concentrations are in micromolar [µM].

**Supplementary Figure 4. VOC content in *Cannabis* essential oils**.

**Supplementary Figure 5. *Anopheles gambiae* OR28 (AgamOR28) does not respond to borneol. a**, Concentration-response of AgamOR28 in response to increasing concentrations of four plant volatile organic compounds, including aromatic (acetophenone and α-terpineol) and terpenoid compounds (α-pinene and (+)-borneol). EC_50_ values shown in the inset are in the low millimolar range. **b**, Representative current traces of AgamOR28-Orco activation by 4 plant volatile organic compounds. Arrowheads above the traces indicate the onset of the odorant stimulus.

**Supplementary Figure 6. Construction of *Ae. aegypti Or49* knockout line. a**, Diagram depicting the *Ae. aegypti Or49* gene and the gRNA target sites on the top and the insertion cassette on the bottom flanked by the upstream (Left) and downstream (Right) homology arms. The insertion cassette encompasses the *QF2* sequence and the 3’ UTR of the *HSP70* gene along with the *ECFP* gene under the control of the *Opie2* promoter and the 3’ UTR of the *SV40* gene. Arrows indicate primer binding sites. Primers P1-12 are listed in Table S4. **b**, Plasmid V1117F-Or49 map. **c**, *In vitro* Cas9 cleavage assay. Guide RNAs left and right targeting the *Or49* gene almost completely digested PCR fragment containing target sequence. (–) negative controls without guide RNAs show no PCR fragment digestion. **d**, Sanger sequencing of amplified PCR fragments of G1 transgenic male individuals unveiling the sequences overlapping upstream (left) and downstream (right) of the DNA cassette inserted by homology directed repair. **e**, Sequencing of G1 individuals showing the complete insertion of the DNA cassette containing *QF2* and *ECFP*. The presence of a single point mutation was confirmed as an artifact and excluded by Sanger sequencing. **f**, Diagnostic PCR reactions showing the presence of double bands in single G1 heterozygous individuals and single upper band in pools of 10 individuals of the *OR49*^-/-^ homozygous line.

**Supplementary Figure 7. The lowest borneol detection threshold of the maxillary palp is 0.1 μg**. Bar plot of the maxillary palp responses to vehicle and increasing doses of borneol. Raw data are shown in the adjacent table. Statistical differences were evaluated via one-way anova. ***p < 0.005 and ****p < 0.001. Data indicate the means ± SEM, *n* = 15-27.

**Table S1. List of blends and single compounds used in the pharmacological screen. Table S2. List of *Or* genes**.

**Table S3. Raw pharmacological data. Table S4. Primer list (*Or49* knockout)**.

## Extended data figures and tables

**Extended Data Fig. 1 Diagrams of the odor delivery system**.

The odor delivery system is composed of interlocking top and bottom rings attached to a removable chemical holder (overview, top, bottom and side views are provided). Dimensions are provided in millimeters.

**Extended Data Fig. 2 Hand rings.stl**.

File format for 3D-printing of the two complementary hand rings.

**Extended Data Fig. 3 Chemical holder.stl**.

File format for 3D-printing of the chemical holder component.

**Extended Data Fig. 4 Behavioral data**.

CSV file of collected mosquito visits on the ROI.

**Extended Data Fig. 5 Video recording example of the of the female *Ae. aegypti* antennal lobe**.

**Extended Data Fig. 6 Video recording example of the arm-in-a-cage assay**.

Thirty-second-long recording example at the 4-minute mark of mosquito landing behavior in the presence of the vehicle diethyl ether (DEE) and racemic borneol.

## Acknowledgments

This study was supported by ISF 997/19 awarded to A.W., ISF 719/21 awarded to J.B., NIH RO1AI175152 and NSF IOS-2242604 awarded to O.S.A., NIH R01AI148300 awarded to J.A.R, O.S.A., and R.J.P. J.A.R was funded by the Bill and Melinda Gates Foundation (INV-021766). Science and Technology Development Plan Project of Jilin Province, China (20200402001NC) awarded to Y.W. This research was supported by the Ministry of Science & Technology, Israel to P.A.P. (grant numbers 3-16795 and 3-17985). We thank Ziv Kassner for consultation, processing, analysis guidance, and technical support in the creation of the mosquito detection model. We are especially grateful to Nadav Eyal at Eybna Terpene Based Technologies for their time and supply of *Cannabis* essential oils. We are grateful to Dr. Laor Orshan (Ministry of Health, Israel) for providing a colony of *Culex pipiens*. Special thanks to Dr. Osnat Malka for productive discussions. Drs. Pitts and Bohbot are co-corresponding authors.

## Contributions

RJP initiated the functional characterization of OR49. JB conceived the study. YV identified borneol as a key OR49 ligand. YW conducted mosquito electropalpograms. RMH identified camphor as an OR49 agonist. OSA and IVC-A engineered the *Ae. aegypti Or49* knockout strain. MG and RI designed and conducted the single sensillum recordings. ESS conducted the behavioral experiments, created the YOLOv8 model and analyzed the data. DP and PP conducted gene annotations, phylogenetic analyses and described OR49 syntenic relationships. DR completed the *Toxorhynchites amboinensis* OR genomic PCRs and sequencing. EY provided genetic constructs and injectable mRNA. AW provided funding and scientific oversight. JB wrote the manuscript with individual contributions from all the authors.

## Corresponding authors

Correspondence to Jonathan D. Bohbot or Ronald J. Pitts.

## Ethics declarations

Competing interests

The authors declare no competing interests.

