## Supplementary Figures 1-7 for "A conserved odorant receptor underpins borneol-mediated repellency in culicine mosquitoes"

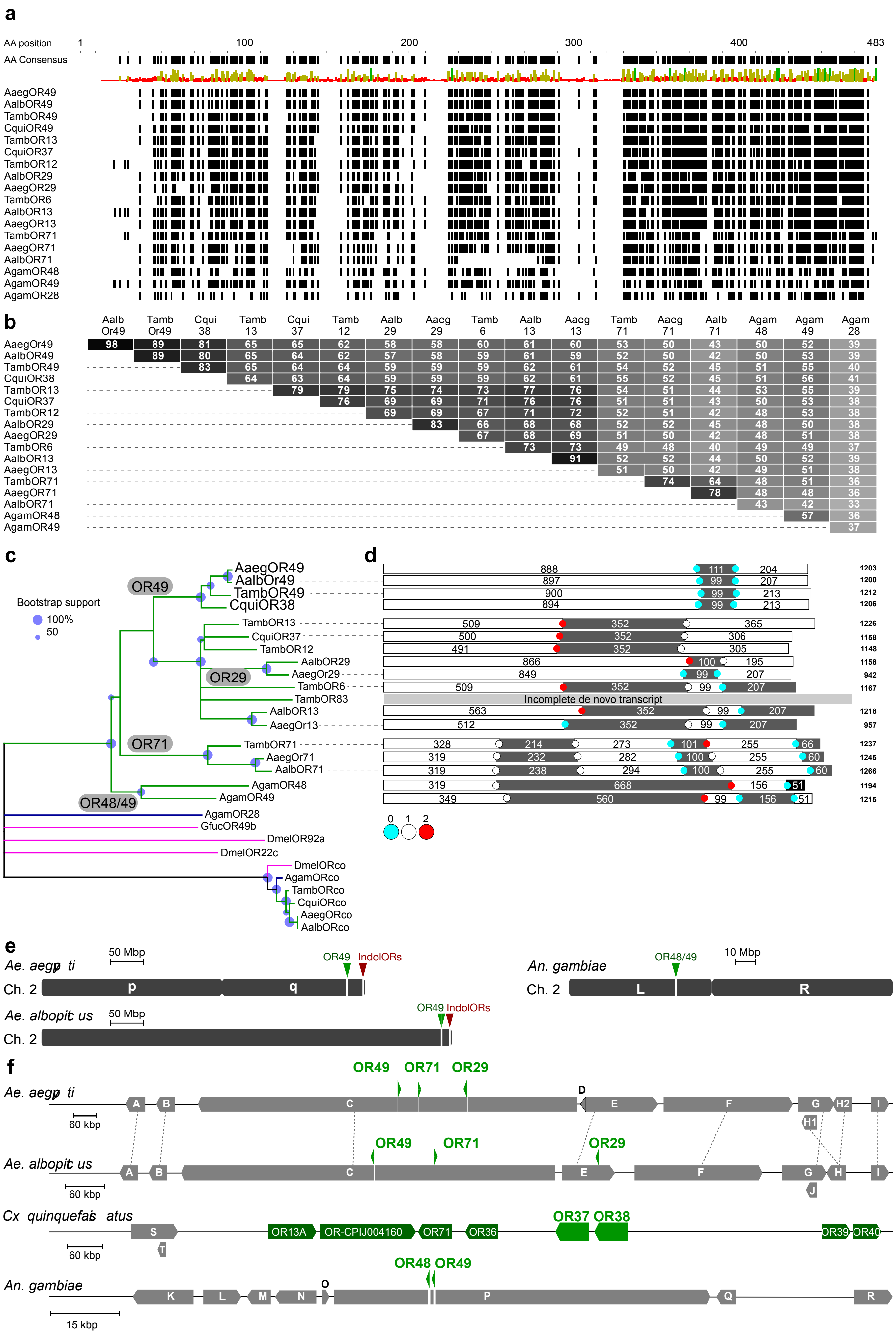
a
AA position
1
100
200
300
400
483
AA Consensus
AaegOR49
AalbOR49
TambOR49
CquiOR49
TambOR13
CquiOR37
TambOR12
AalbOR29
AaegOR29
TambOR6
AalbOR13
AaegOR13
TambOR71
AaegOR71
AalbOR71
AgamOR48
AgamOR49
AgamOR28
b
Aalb Or49 | Tamb Or49 | Cqui 38 | Tamb 13 | Cqui 37 | Tamb 12 | Aalb 29 | Aaeg 29 | Tamb 6 | Aalb 13 | Aaeg 13 | Tamb 71 | Aaeg 71 | Aalb 71 | Agam 48 | Agam 49 | Agam 28
AaegOr49 98 89 81 65 65 62 58 58 60 61 60 53 50 43 50 52 39
AalbOR49 89 80 65 64 62 57 58 59 61 59 52 50 42 50 53 39
TambOR49 83 65 64 64 59 59 59 62 61 54 52 45 51 55 40
CquiOR38 64 63 64 59 59 59 62 61 55 52 45 51 56 41
TambOR13 79 79 75 74 73 77 76 54 51 44 53 55 39
CquiOR37 76 69 69 71 76 76 51 51 43 50 53 38
TambOR12 69 69 67 71 72 52 51 42 48 53 38
AalbOR29 83 66 68 68 52 52 45 48 50 38
AaegOR29 67 68 69 51 50 42 48 51 38
TambOR6 73 73 49 48 40 49 49 37
AalbOR13 91 52 52 44 50 52 39
AaegOR13 51 50 42 49 51 38
TambOR71 74 64 48 51 36
AaegOR71 78 48 48 36
AalbOR71 43 42 33
AgamOR48 57 36
AgamOR49 37
c
Bootstrap support
100%
50
OR49
OR29
OR71
OR48/49
AaegOR49
AalbOr49
TambOR49
CquiOR38
TambOR13
CquiOR37
TambOR12
AalbOR29
AaegOr29
TambOR6
TambOR83
AalbOR13
AaegOr13
TambOR71
AaegOr71
AalbOR71
AgamOR48
AgamOR49
AgamOR28
GfucOR49b
DmelOR92a
DmelOR22c
DmelORco
AgamORco
TambORco
CquiORco
AaegORco
AalbORco
d
888 111 204 1203
897 99 207 1200
900 99 213 1212
894 99 213 1206
509 352 365 1226
500 352 306 1158
491 352 305 1148
866 100 195 1158
849 99 207 942
509 352 99 207 1167
Incomplete de novo transcript
563 352 99 207 1218
512 352 99 207 957
328 214 273 101 255 66 1237
319 232 282 100 255 60 1245
319 238 294 100 255 60 1266
319 668 156 51 1194
349 560 99 156 51 1215
0 1 2
e
Ae. aegypti
50 Mbp
OR49
IndolORs
Ch. 2
p
q
An. gambiae
OR48/49
10 Mbp
Ch. 2
L
R
Ae. albopictus
50 Mbp
OR49 IndolORs
Ch. 2
f
OR49 OR71 OR29
Ae. aegypti
D
A B C E F G H2 I
H1
60 kbp
OR49 OR71 OR29
Ae. albopictus
A B C E F G H I
J
60 kbp
Cx. quinquefasciatus
S
T
OR13A OR-CPIJ004160 OR71 OR36
OR37 OR38
OR39 OR40
60 kbp
OR48 OR49
An. gambiae
O
K L M N P Q R
15 kbp

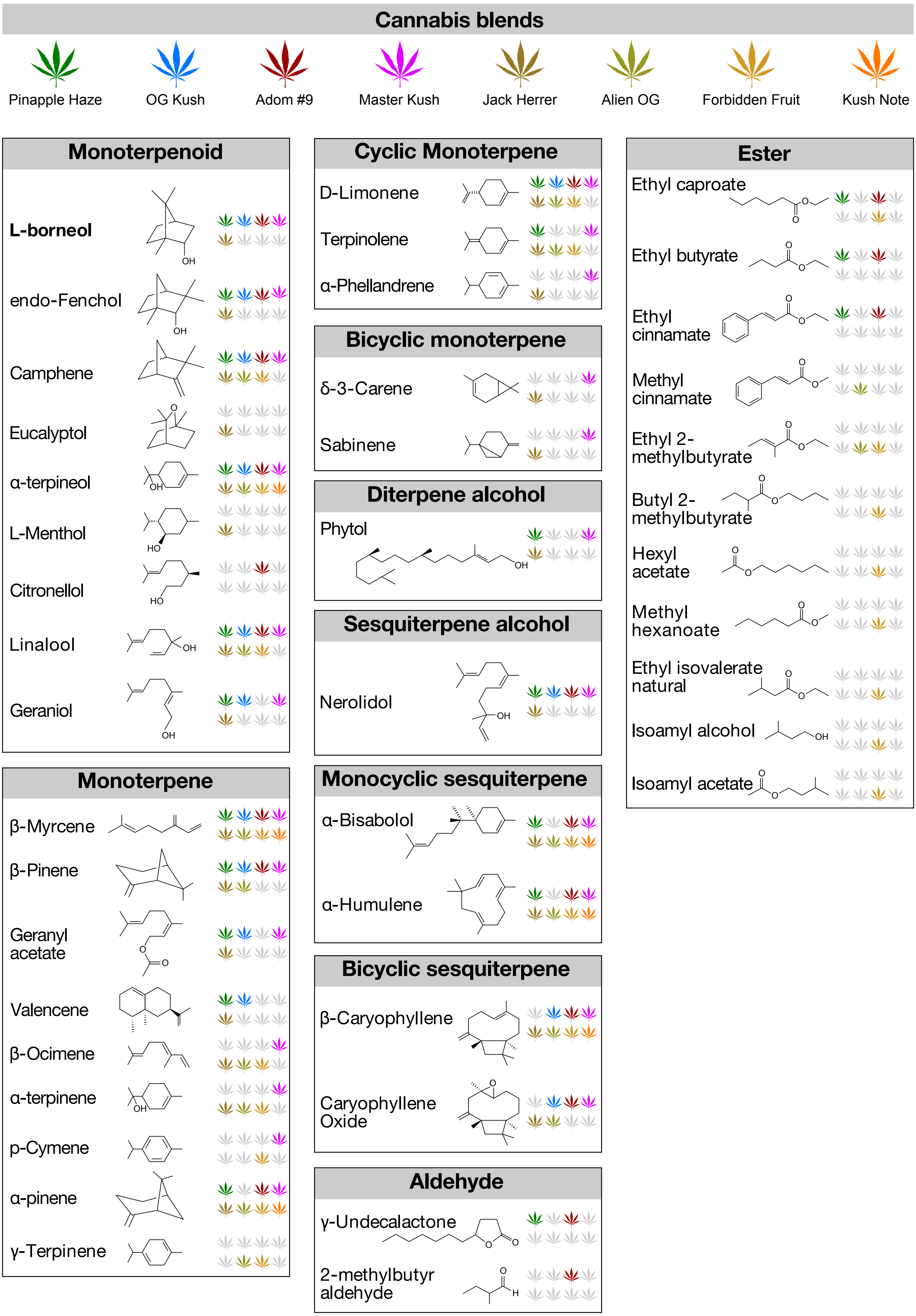
Cannabis blends
Pinapple Haze
OG Kush
Adom #9
Master Kush
Jack Herrer
Alien OG
Forbidden Fruit
Kush Note
Monoterpenoid
L-borneol
endo-Fenchol
Camphene
Eucalyptol
α-terpineol
L-Menthol
Citronellol
Linalool
Geraniol
Monoterpene
β-Myrcene
β-Pinene
Geranyl acetate
Valencene
β-Ocimene
α-terpinene
p-Cymene
α-pinene
γ-Terpinene
Cyclic Monoterpene
D-Limonene
Terpinolene
α-Phellandrene
Bicyclic monoterpene
δ-3-Carene
Sabinene
Diterpene alcohol
Phytol
Sesquiterpene alcohol
Nerolidol
Monocyclic sesquiterpene
α-Bisabolol
α-Humulene
Bicyclic sesquiterpene
β-Caryophyllene
Caryophyllene Oxide
Aldehyde
γ-Undecalactone
2-methylbutyr aldehyde
Ester
Ethyl caproate
Ethyl butyrate
Ethyl cinnamate
Methyl cinnamate
Ethyl 2-methylbutyrate
Butyl 2-methylbutyrate
Hexyl acetate
Methyl hexanoate
Ethyl isovalerate natural
Isoamyl alcohol
Isoamyl acetate

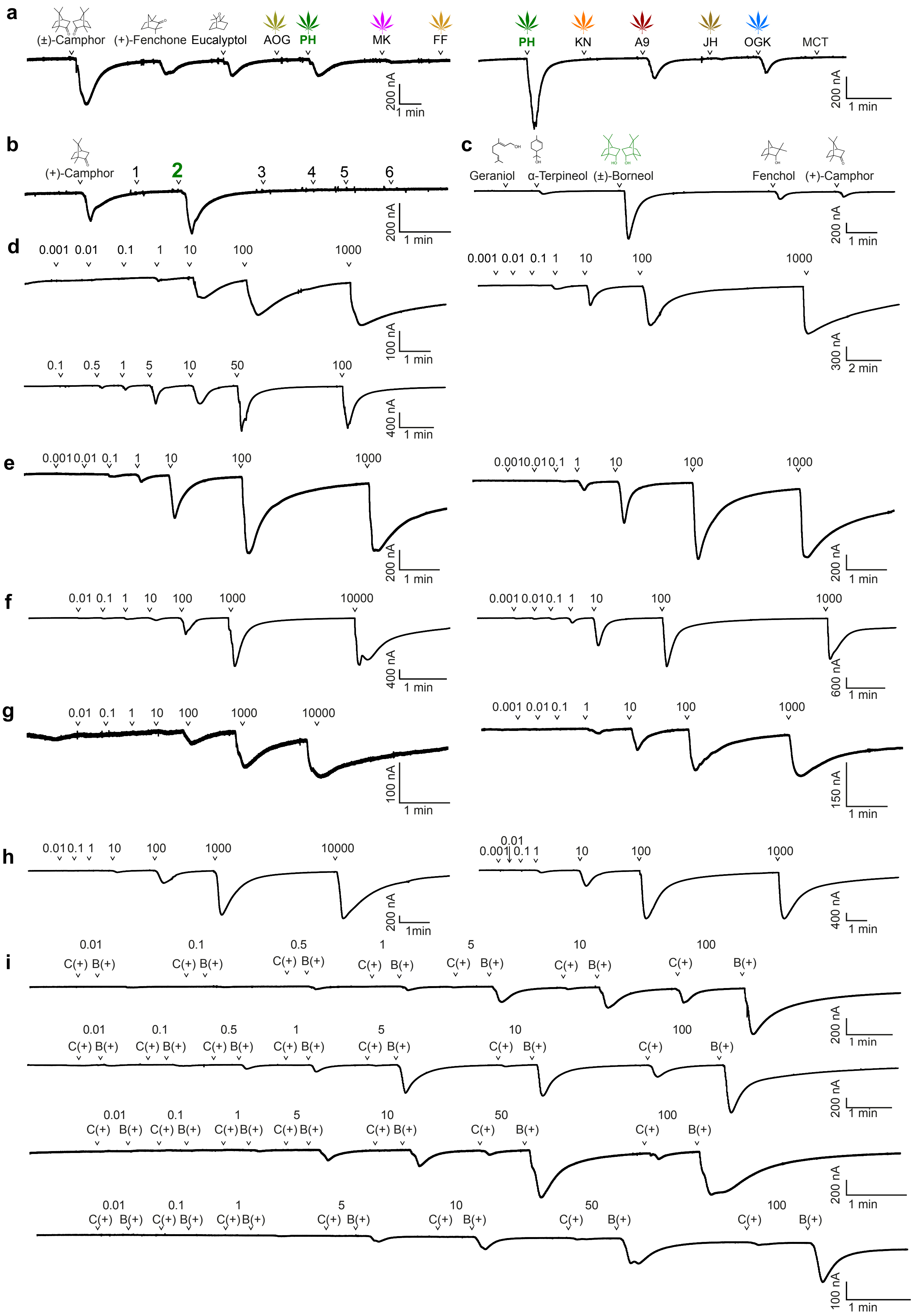
a
(±)-Camphor
(+)-Fenchone
Eucalyptol
AOG
PH
MK
FF
200 nA
1 min
PH
KN
A9
JH
OGK
MCT
200 nA
1 min
b
(+)-Camphor
1
2
3
4
5
6
200 nA
1 min
c
Geraniol
α-Terpineol
(±)-Borneol
Fenchol
(+)-Camphor
200 nA
1 min
d
0.001 0.01 0.1 1 10 100 1000
100 nA
1 min
0.001 0.01 0.1 1 10 100 1000
300 nA
2 min
0.1 0.5 1 5 10 50 100
400 nA
1 min
e
0.001 0.01 0.1 1 10 100 1000
200 nA
1 min
0.001 0.01 0.1 1 10 100 1000
200 nA
1 min
f
0.01 0.1 1 10 100 1000 10000
400 nA
1 min
0.001 0.01 0.1 1 10 100 1000
600 nA
1 min
g
0.01 0.1 1 10 100 1000 10000
100 nA
1 min
0.001 0.01 0.1 1 10 100 1000
150 nA
1 min
h
0.01 0.1 1 10 100 1000 10000
200 nA
1min
0.01
0.001 0.1 1 10 100 1000
400 nA
1 min
i
0.01 0.1 0.5 1 5 10 100
C(+) B(+)
200 nA
1 min
0.01 0.1 0.5 1 5 10 100
C(+) B(+)
200 nA
1 min
0.01 0.1 1 5 10 50 100
C(+) B(+)
200 nA
1 min
0.01 0.1 1 5 10 50 100
C(+) B(+)
100 nA
1 min

**a**

| Pineapple Haze | |
|---|---|
| VOC | Weight % |
| β-Myrcene | 17.42 |
| D-Limonene | 10.8 |
| Linalool | 9.77 |
| α-Bisabolol | 8.85 |
| α-Humulene | 7.62 |
| β-Pinene | 3.03 |
| Phytol | 2.39 |
| endo-Fenchol | 2.03 |
| α-Pinene | 1.99 |
| Nerolidol | 1.83 |
| α-Terpineol | 1.64 |
| Ethyl caproate | 1.45 |
| **(L)-(–)-borneol** | **1.32** |
| Geranyl Acetate | 1.12 |
| Geraniol | 1 |
| Camphene | 1 |
| Terpinolene | 0.84 |
| Ethyl Butyrate | 0.22 |
| Ethyl Cinnamate | 0.15 |
| Valencene | 0.12 |
| γ-Undecalactone | 0.07 |

| OG Kush | |
|---|---|
| VOC | Weight % |
| β-Myrcene | 32.66 |
| D-Limonene | 24.93 |
| β-Caryophyllene | 21.17 |
| Linalool | 5.97 |
| β-Pinene | 3.61 |
| Nerolidol | 3.01 |
| α-Terpineol | 2.76 |
| endo-Fenchol | 2.45 |
| Geranyl Acetate | 1.49 |
| **(L)-(–)-borneol** | **0.88** |
| Caryophyllene Oxide | 0.39 |
| Valencene | 0.37 |
| Camphene | 0.22 |
| Geraniol | 0.09 |

| Adom #9 | |
|---|---|
| VOC | Weight % |
| β-Myrcene | 29.12 |
| β-Caryophyllene | 23.39 |
| α-Pinene | 11.62 |
| α-Humulene | 7.82 |
| β-Pinene | 6.46 |
| Linalool | 5.44 |
| D-Limonene | 4.99 |
| α-Bisabolol | 2.98 |
| Nerolidol | 2.41 |
| α-Terpineol | 1.77 |
| endo-Fenchol | 1.32 |
| Caryophyllene Oxide | 1.18 |
| **(L)-(–)-borneol** | **0.58** |
| Citronellol | 0.42 |
| Camphene | 0.28 |
| Fenchone | 0.22 |
| Ethyl caproate | 0.0073 |
| Ethyl Butyrate | 0.0011 |
| 2-methylbutyraldehyde | 0.001 |
| Ethyl Cinnamate | 0.0007 |
| γ-Undecalactone | 0.0004 |

| Master Kush | |
|---|---|
| VOC | Weight % |
| β-Myrcene | 46.39 |
| β-Caryophyllene | 8.28 |
| D-Limonene | 6.16 |
| β-Ocimene | 4.7 |
| α-Terpineol | 3.04 |
| Phytol | 2.95 |
| Nerolidol | 2.7 |
| α-Phellandrene | 2.37 |
| δ-3-Carene | 2.37 |
| α-Terpinene | 2.37 |
| Sabinene | 2.37 |
| p-cymene | 2.37 |
| Geranyl Acetate | 2.37 |
| α-Humulene | 2.33 |
| Linalool | 2.29 |
| β-Pinene | 1.5 |
| endo-Fenchol | 0.79 |
| α-Pinene | 0.71 |
| **(L)-(–)-borneol** | **0.33** |
| Caryophyllene Oxide | 0.33 |
| Camphene | 0.17 |
| Terpinolene | 0.04 |
| Geraniol | 0.04 |
| α-Bisabolol | 0.04 |

| Jack Herer | |
|---|---|
| VOC | Weight % |
| Terpinolene | 39.41 |
| β-Caryophyllene | 14.81 |
| β-Ocimene | 10.22 |
| β-Myrcene | 7.04 |
| D-Limonene | 5.09 |
| Phytol | 4.5 |
| β-Pinene | 3.87 |
| Nerolidol | 2.69 |
| Alpha-Phellandrene | 2.5 |
| δ-3-Carene | 1.6 |
| α-Pinene | 1.48 |
| α-Terpineol | 1.46 |
| α-Humulene | 1.39 |
| α-Terpinene | 1.37 |
| Caryophyllene Oxide | 0.72 |
| Linalool | 0.57 |
| α-Bisabolol | 0.39 |
| endo-Fenchol | 0.26 |
| Geranyl Acetate | 0.15 |
| Eucalyptol | 0.12 |
| Sabinene | 0.11 |
| **(L)-(–)-borneol** | **0.07** |
| Geraniol | 0.07 |
| Camphene | 0.07 |
| L-menthol | 0.027 |
| Valencene | 0.01 |

| Alien OG | |
|---|---|
| VOC | Weight % |
| β-Caryophyllene | 25.69 |
| β-Myrcene | 23.59 |
| D-Limonene | 17.27 |
| Terpinolene | 10.79 |
| Linalool | 6.782 |
| α-Pinene | 4.134 |
| α-Bisabolol | 3.516 |
| β-Ocimene | 3.391 |
| β-Pinene | 2.47 |
| Camphene | 0.6174 |
| α-Terpinene | 0.6174 |
| Caryophyllene Oxide | 0.6174 |
| γ-Terpinene | 0.304 |
| α-Terpineol | 0.125 |
| α-Humulene | 0.125 |
| Ethyl 2-methylbutyrate | 0.02 |
| Methyl cinnamate | 0.01 |

| Forbidden Fruit | |
|---|---|
| VOC | Weight % |
| β-Myrcene | 26.97 |
| D-Limonene | 19.39 |
| β-Caryophyllene | 15.28 |
| Terpinolene | 14.71 |
| Linalool | 6.347 |
| α-Pinene | 5.398 |
| β-Ocimene | 4.682 |
| β-Pinene | 2.008 |
| α-Bisabolol | 1.725 |
| α-Terpinene | 1.342 |
| Camphene | 0.9991 |
| Gamma-Terpinene | 0.6661 |
| p-cymene | 0.333 |
| α-Terpineol | 0.05 |
| α-Humulene | 0.05 |
| Ethyl 2-methylbutyrate | 0.0117 |
| Ethyl caproate | 0.00877 |
| Butyl 2-methylbutyrate | 0.00877 |
| Hexyl acetate | 0.005845 |
| Methyl hexanoate | 0.004675 |
| Ethyl isovalerate natural | 0.004385 |
| Isoamyl alcohol | 0.00292 |
| Isoamyl acetate | 0.00292 |

| Kush Note | |
|---|---|
| VOC | Weight % |
| β-Caryophyllene | 44.91 |
| β-Myrcene | 44.91 |
| α-Terpineol | 2.5 |
| α-Humulene | 2.5 |
| α-Bisabolol | 2.5 |
| α-Pinene | 2.5 |

**b**

| Sub-mixture 1 | |
|---|---|
| VOC | Weight % |
| Myrcene | 33.33 |
| Limonene | 33.33 |
| Linalool | 33.33 |

| Sub-mixture 2 | |
|---|---|
| VOC | Weight % |
| Fenchol | 25 |
| **(±)--Borneol** | **25** |
| Terpineol | 25 |
| Geraniol | 25 |

| Sub-mixture 3 | |
|---|---|
| VOC | Weight % |
| β-Caryophyllene | 25 |
| Caryophyllene oxide | 25 |
| Camphene | 25 |
| β-Pinene | 25 |

| Sub-mixture 4 | |
|---|---|
| VOC | Weight % |
| Nerolidol | 33.33 |
| Geranyl acetate | 33.33 |
| Valencene | 33.33 |

| Sub-mixture 5 | |
|---|---|
| VOC | Weight % |
| Ethyl Butyrate | 50 |
| Ethyl caproate | 50 |

| Sub-mixture 6 | |
|---|---|
| VOC | Weight % |
| Ethyl Cinnamate | 50 |
| γ-Undecalactone | 50 |

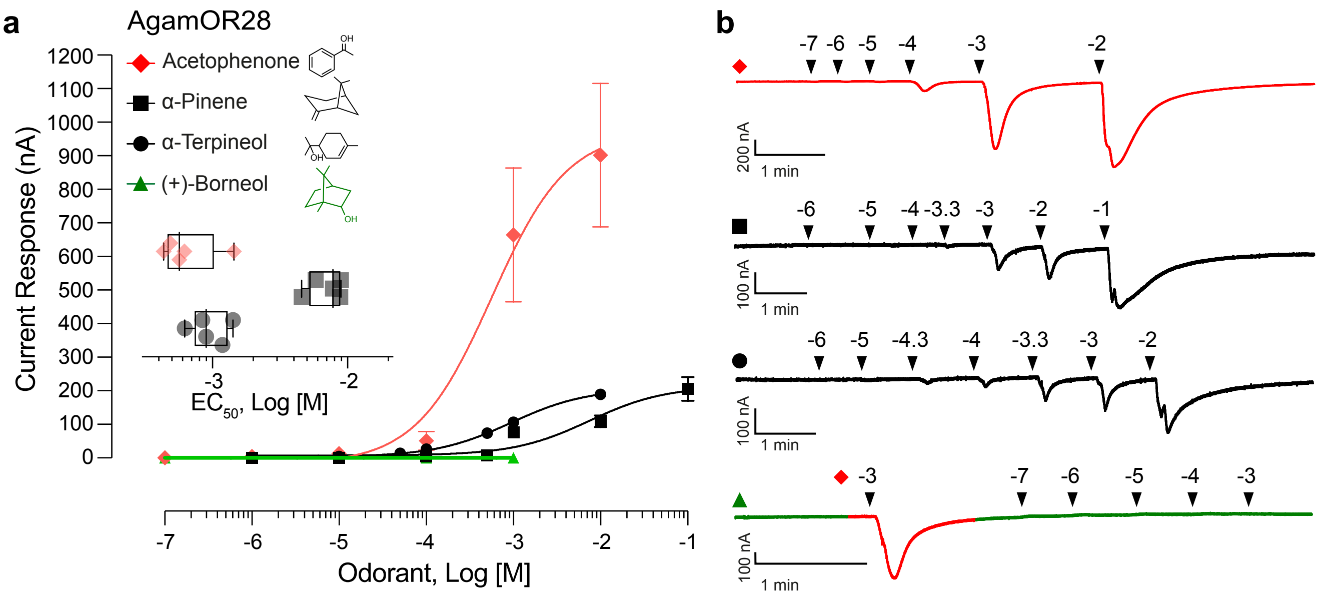
a
AgamOR28
Acetophenone
α-Pinene
α-Terpineol
(+)-Borneol
Current Response (nA)
EC50, Log [M]
Odorant, Log [M]
b
200 nA
1 min
100 nA
1 min
100 nA
1 min
100 nA
1 min

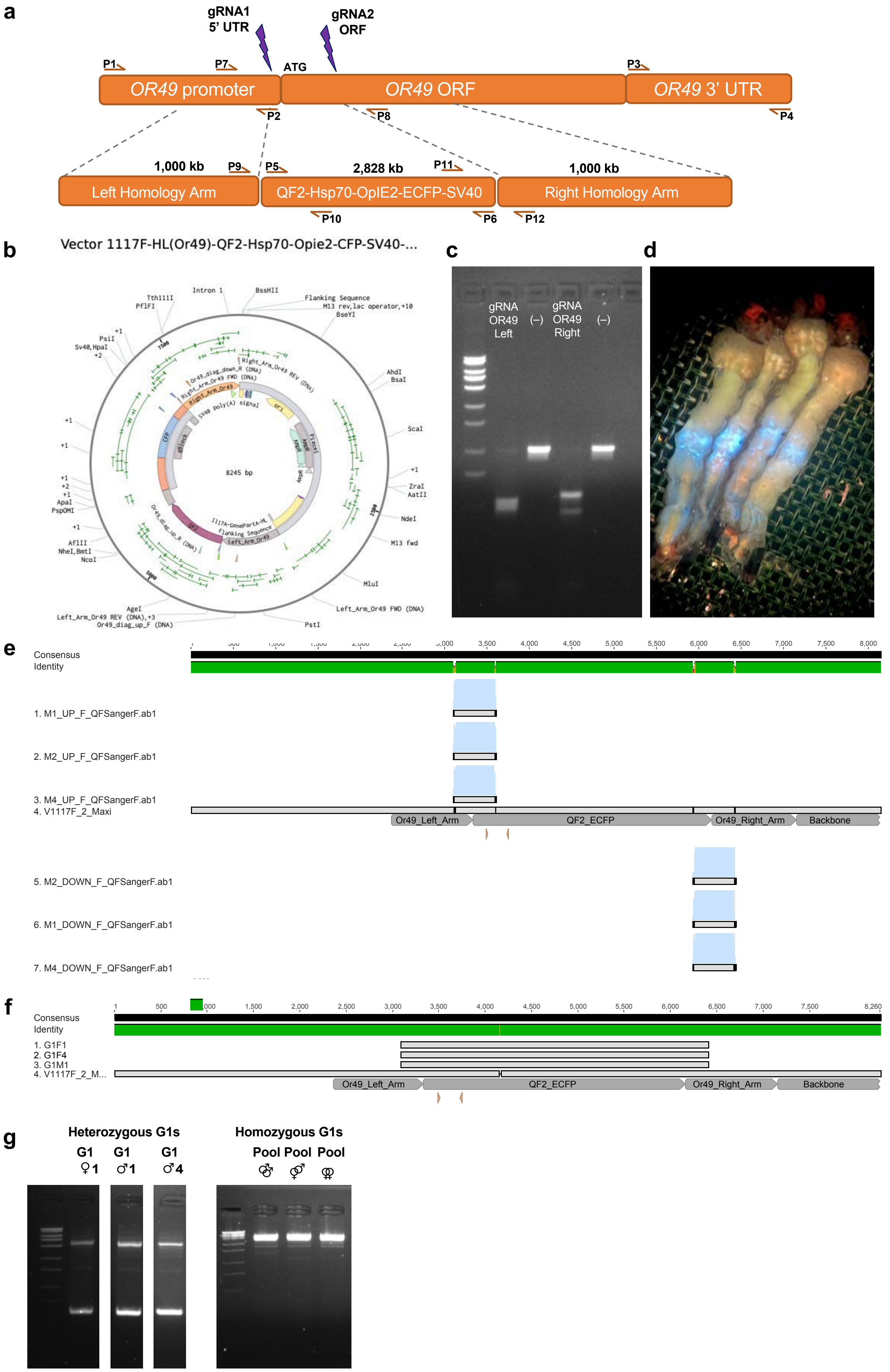
a
gRNA1
5' UTR
gRNA2
ORF
ATG
P1
P7
P3
OR49 promoter
OR49 ORF
OR49 3' UTR
P2
P8
P4
1,000 kb
P9
P5
2,828 kb
P11
1,000 kb
Left Homology Arm
QF2-Hsp70-OpIE2-ECFP-SV40
Right Homology Arm
P10
P6
P12
b
Vector 1117F-HL(Or49)-QF2-Hsp70-Opie2-CFP-SV40-...
8245 bp
c
gRNA OR49 Left
(–)
gRNA OR49 Right
(–)
d
e
Consensus
Identity
1. M1_UP_F_QFSangerF.ab1
2. M2_UP_F_QFSangerF.ab1
3. M4_UP_F_QFSangerF.ab1
4. V1117F_2_Maxi
Or49_Left_Arm
QF2_ECFP
Or49_Right_Arm
Backbone
5. M2_DOWN_F_QFSangerF.ab1
6. M1_DOWN_F_QFSangerF.ab1
7. M4_DOWN_F_QFSangerF.ab1
f
Consensus
Identity
1. G1F1
2. G1F4
3. G1M1
4. V1117F_2_M...
Or49_Left_Arm
QF2_ECFP
Or49_Right_Arm
Backbone
g
Heterozygous G1s
G1 ♀1
G1 ♂1
G1 ♂4
Homozygous G1s
Pool
Pool
Pool

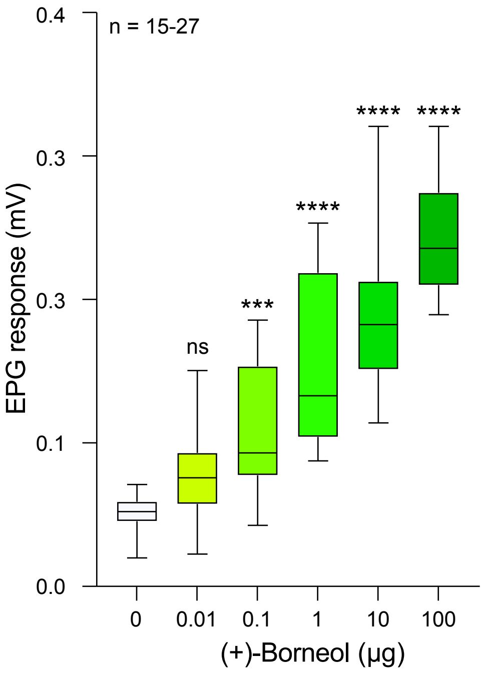


| 0 µg | 0.01 µg | 0.1 µg | 1 µg | 10 µg | 100 µg |
|---|---|---|---|---|---|
| 0.04544 | 0.14768 | 0.16472 | 0.21868 | 0.19596 | 0.228904 |
| 0.03124 | 0.15052 | 0.15336 | 0.17892 | 0.1988 | 0.23856 |
| 0.04544 | 0.14768 | 0.13632 | 0.1846 | 0.20164 | 0.23572 |
| 0.0568 | 0.054868 | 0.185707 | 0.244795 | 0.320766 | 0.27856 |
| 0.06248 | 0.084412 | 0.151942 | 0.253237 | 0.312325 | 0.27434 |
| 0.06248 | 0.059089 | 0.160383 | 0.219472 | 0.320766 | 0.320766 |
| 0.059089 | 0.07972 | 0.043182 | 0.132867 | 0.166083 | 0.255768 |
| 0.050647 | 0.056468 | 0.093007 | 0.102972 | 0.212586 | 0.288985 |
| 0.042206 | 0.093007 | 0.096328 | 0.112937 | 0.113933 | 0.25909 |
| 0.05979 | 0.071001 | 0.080468 | 0.108868 | 0.151469 | 0.189336 |
| 0.053147 | 0.075735 | 0.080468 | 0.094668 | 0.156203 | 0.19407 |
| 0.05979 | 0.075735 | 0.085201 | 0.104135 | 0.151469 | 0.198803 |
| 0.051486 | 0.022512 | 0.042523 | 0.087547 | 0.147579 | 0.227622 |
| 0.033217 | 0.057531 | 0.05503 | 0.122566 | 0.155083 | 0.210113 |
| 0.046503 | 0.067536 | 0.077542 | 0.150081 | 0.182598 | 0.225121 |
| 0.01988 | | | | | |
| 0.071001 | | | | | |
| 0.061534 | | | | | |
| 0.047334 | | | | | |
| 0.052068 | | | | | |
| 0.052068 | | | | | |
| 0.050027 | | | | | |
| 0.05503 | | | | | |
| 0.053779 | | | | | |
| 0.047526 | | | | | |
| 0.057531 | | | | | |
| 0.040022 | | | | | |
