## Supplementary figures and images for "A conserved odorant receptor underpins borneol-mediated repellency in culicine mosquitoes"

### Extended Figure 1

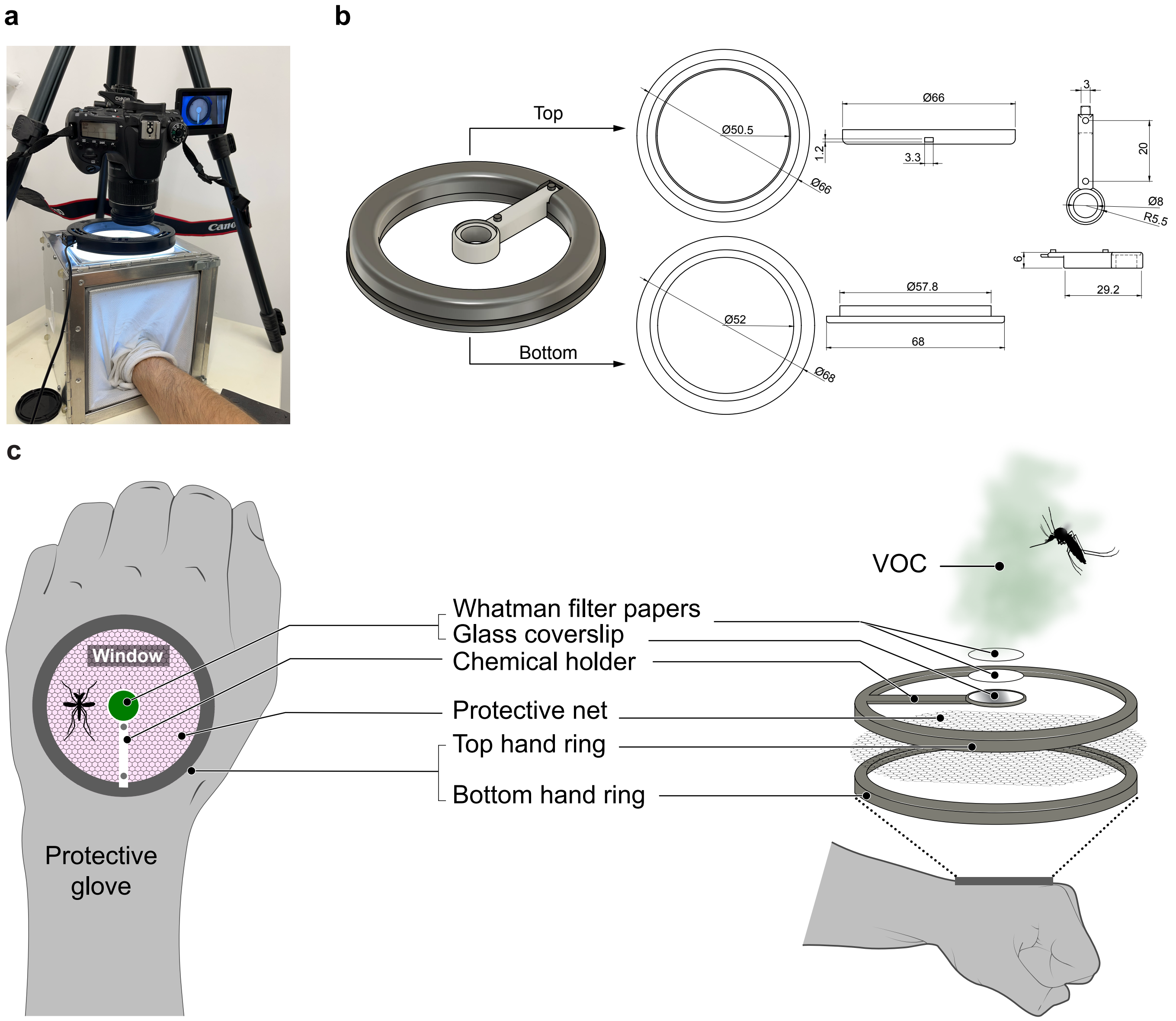
